# Three-dimensional spatial transcriptomics at isotropic resolution enabled by generative deep learning

**DOI:** 10.1101/2025.08.15.670472

**Authors:** Bohan Li, Feng Bao, Yuhang Sun, Tianyun Wang, Fengji Li, Hongjue Li, Qionghai Dai, Yue Deng

**Affiliations:** School of Astronautics, Beihang University, Beijing 102206, China; College of Future Information Technology, Fudan University, Shanghai 200433, China; Institute of Artificial Intelligence, Beihang University, Beijing 100191, China; Institute for Brain and Cognitive Sciences, Tsinghua University, Beijing 100084, China

## Abstract

Mapping the complete three-dimensional (3D), transcriptome-wide spatial architecture of tissues and organs remains a fundamental challenge in biology. Typically, studies approximate 3D structures by profiling serial two-dimensional (2D) tissue sections, resulting in incomplete and anisotropic reconstructions. Here, we demonstrate that organ-wide, isotropic-resolution 3D gene expression landscapes can be effectively inferred from sparse 2D spatial transcriptomics data using artificial intelligence. Our framework, isoST, leverages stochastic differential equations (SDEs) to model the intrinsic biological continuity of gene expression along tissue depth, enabling the integration of discrete 2D measurements into coherent 3D transcriptomic maps. isoST is applied across diverse biological systems, including the brain, kidney, spinal cord, and gastrulating embryos, accurately reconstructing anatomical structures and molecular patterns that are only partially resolved in 2D analyses. isoST enables the comprehensive characterization of tissue niches at full 3D resolution, uncovering radial and spherical gene expression gradients associated with disease progression that are often distorted or missed in 2D approaches. Furthermore, isoST is extended to integrate histological imaging for reducing experimental complexity and to reconstruct high-resolution developmental spatial trajectories over time. Together, isoST provides a practical and scalable solution toward comprehensive high-resolution 3D spatial transcriptomic atlases.

## Introduction

Resolving the spatial context of biomolecules within intact tissues is fundamental to understanding biological processes and disease mechanisms(*1-3*). Recent spatial omics technologies have significantly enhanced our capacity for spatially resolved transcriptional profiling, yet current platforms remain largely constrained to two-dimensional (2D) tissue sections(*4-6*). However, biological tissues are inherently three-dimensional (3D) architectures, and many essential processes, including cell-cell interactions across tissue depths(*7, 8*), neural circuit formation in brain layers(*7, 9-12*), and complex tumor microenvironment organization(*13, 14*), can only be fully understood within a true 3D context. Accurately mapping isotropic-resolution 3D spatial transcriptomic profiles therefore represents a fundamental challenge in biology.

Several recent technologies have attempted to address this challenge. TRISCO(*15*) applies enhanced tissue-clearing protocols to image multiple RNA species throughout entire organs; Deep-STARmap(*16*) integrates cDNA crosslinking with hydrogel embedding to visualize RNAs within substantial tissue volumes; and cycleHCR(*17*) utilizes iterative DNA barcoding with hybridization chain reaction to profile 3D RNA distributions in whole embryos. While powerful, these approaches require elaborate workflows with sophisticated instrumentation, extensive processing, and highly specialized expertise, constraining their scalability and widespread adoption. Additionally, their dependence on selective markers and imaging strategies limits transcriptome-wide coverage.

Given the current technological limitations in directly achieving comprehensive 3D transcriptomic profiling, a practical compromise is to section tissues into multiple serial 2D slices along the depth axis and profile each slice individually using commercially available spatial omics platforms(*18-24*). Although this approach provides high-resolution, transcriptomics-wide data within each 2D plane (typically thousands to millions of spots or cells per slice, the depth dimension is typically sampled sparsely (often limited to several to tens of sections)(*18-24*). This imbalance produces coarse, discontinuous, and anisotropic representations of the underlying 3D tissue architecture.

We hypothesize that continuous, isotropic-resolution 3D transcriptomic landscapes can be reconstructed from existing serial 2D spatial omics datasets. However, this task poses significant challenges. Standard spatial regression models are prone to overfitting along the sparsely sampled depth axis, where the number measurements are often several orders of magnitude lower than within individual planes. Meanwhile, adjacent tissue sections may exhibit abrupt molecular or structural differences, making simple interpolation approaches insufficient.

Here, we developed isoST to reconstruct 3D spatial transcriptomic profiles with isotropic resolutions from sparsely sampled serial sections. The key principle behind isoST is that gene expression varies smoothly across biological tissues in 3D space. isoST captures this spatial continuity by using stochastic differential equations (SDEs) to model gene expression dynamics along the tissue depth. Training on profiles from serial tissue sections, isoST defines a continuous 3D spatial transcriptomic field, allowing for the reconstruction of the full spatial continuum at isotropic resolution.

We evaluate isoST across diverse biological systems in both healthy and diseased states, including mouse brain, kidney, and spinal cord, demonstrating its ability to accurately reconstruct organ-wide complex transcriptomic structures from sparsely sampled sections. We show the resulting 3D spatial transcriptomic continuum reveals fine-scale tissue architecture and spatial disease patterns that are not discernible in 2D, enabling downstream analyses tailored to volumetric data, such as domain segmentation and 3D gene expression gradient mapping. By integrating serial two-photon tomography (STPT) images(*25*) as structural priors, isoST maintains high reconstruction quality while reducing the number of sections required. Finally, we extend isoST to temporal contexts, capturing continuous developmental trajectories and periodic gene expression dynamics during mouse embryogenesis.

## Results

### isoST architecture

Given a set of *K* serial 2D spatial transcriptomic sections at discrete depths 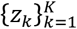, isoST reconstructs an isotropic 3D gene expression continuum over the full tissue depth *z* ∈ [*z*_1_, *z*_*K*_] (**Figs. 1A, B**; **Movie S1**). isoST models intrinsic spatial continuity of biological tissues along the z-axis using two stochastic differential equations (SDEs): a shape SDE that describes tissue structure deformation, and an expression SDE that describes the evolution of gene expression levels across depth (**Fig. 1C**; **Supplementary Fig. 1A**). In each SDE, a trainable gradient function is used to learn the shape or expression change at depth *z* (**Methods**). Starting from an observed profile at depth *z*_*k*_, the expression at a nearby unprofiled depth *z*_*k*_ + Δ*z* is predicted as a smooth continuation guided by the learned gradients. By iteratively applying this procedure across slices with small steps, isoST constructs near-continuous gene expression transitions along the *z*-axis, yielding an isotropic 3D spatial transcriptomic continuum (**Fig. 1B**; **Supplementary Fig. 1B**; **Methods**).

**Figure 1.**
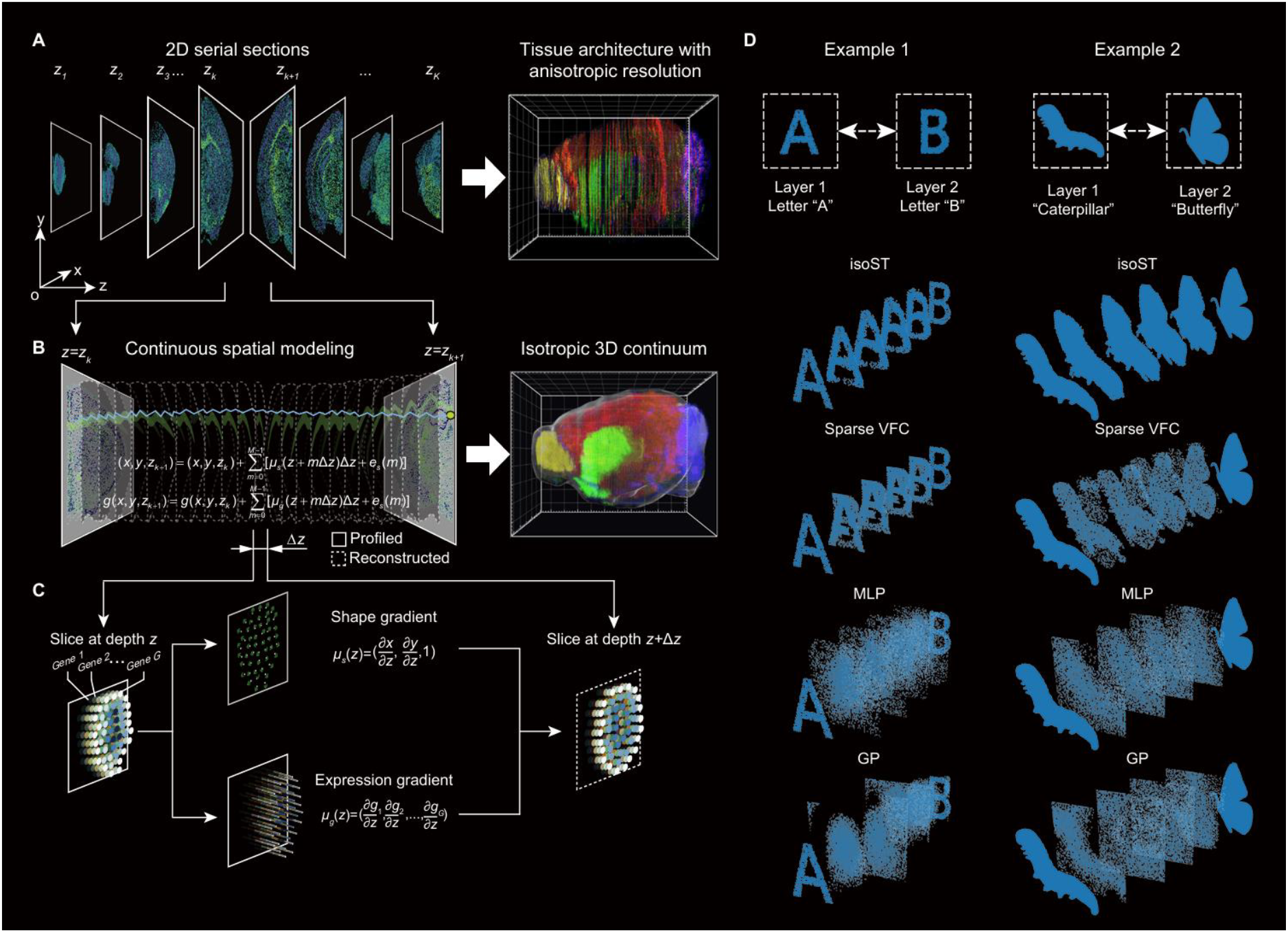
Reconstruction of 3D spatial transcriptomics at isotropic resolution. (**A**) isoST takes as input a series of *K* parallel two-dimensional (2D) spatial transcriptomics slices at discrete depths *z*_1_, *z*_2_, …, *z*_*K*_. (**B**) isoST models spatial continuity along the *z* -axis using stochastic differential equations (SDEs) to reconstruct 3D transcriptomics profiles at isotropic resolution. Starting from an observed slice at depth *z*_*k*_, the model iteratively propagates each cell’s spatial position (*x, y, z*_*k*_) and gene expression ***g***(*x, y, z*_*k*_) to the next layer (*x*’, *y*’, *z*_*k*+1_) through integration over *M* small steps of size Δ*z*. (**C**) A schematic of a single reconstruction step from depth *z* to *z* + Δ*z*. The shape gradient term ***μ***_s_(*z*) determines the directional shift in position for each cell, while the expression gradient term ***μ***_g_(*z*) estimates the gradient of gene expression used to impute the next layer. (**D**) Toy examples demonstrating isoST’s ability to model smooth transitions between sparsely sampled layers. In example 1 (left), a morphing sequence from letter “A” to “B”; in example 2 (right), a transformation from a caterpillar to a butterfly.

### Demonstration using synthetic data

To conceptually evaluate the effectiveness of isoST, we designed synthetic benchmarks consisting of only two spatial layers, each representing distinct 2D shapes (for example, the letters “A” and “B” or forms such as a caterpillar and a butterfly). The goal was to infer a smooth, spatially coherent transition between the two layers. isoST successfully reconstructed continuous transformations that preserved both the overall shape and internal structure (**Fig. 1D**). In contrast, the interpolation-based method (Sparse VFC(*26*)) yielded blurred intermediates that resembled simple mixtures of the input shapes, while position-based regressors such as the multilayer perceptron (MLP) and the Gaussian process (GP) failed to produce coherent results due to the large differences between the two input layers (**Methods**). These findings demonstrate isoST’s ability to learn spatial transitions from minimal input, validating its core framework for 3D reconstruction under sparse conditions.

### isoST reconstructs isotropic-resolution 3D spatial transcriptomes of the mouse brain

As an initial test case, we applied isoST to a MERFISH-based(*5*) mouse brain dataset consisting of 54 coronal sections at 0.2 mm intervals, each profiling 1,122 genes(*19*) (**Fig. 2A**, top; **Methods**). To assess reconstruction accuracy, isoST was trained on 27 slices, and the inferred 3D transcriptomic field was evaluated against the 26 held-out slices across all genes (**Supplementary Fig. 2A)**. isoST outperformed interpolation-based (Sparse VFC)(*26*) and regression-based (MLP, GP) methods, achieving lower mean squared error (MSE) and higher Dice coefficients (**Supplementary Fig. 2B**). The reconstructed slices also closely matched the ground truth in cell type composition and spatial organization (**Supplementary Figs. 2C, D**).

**Figure 2.**
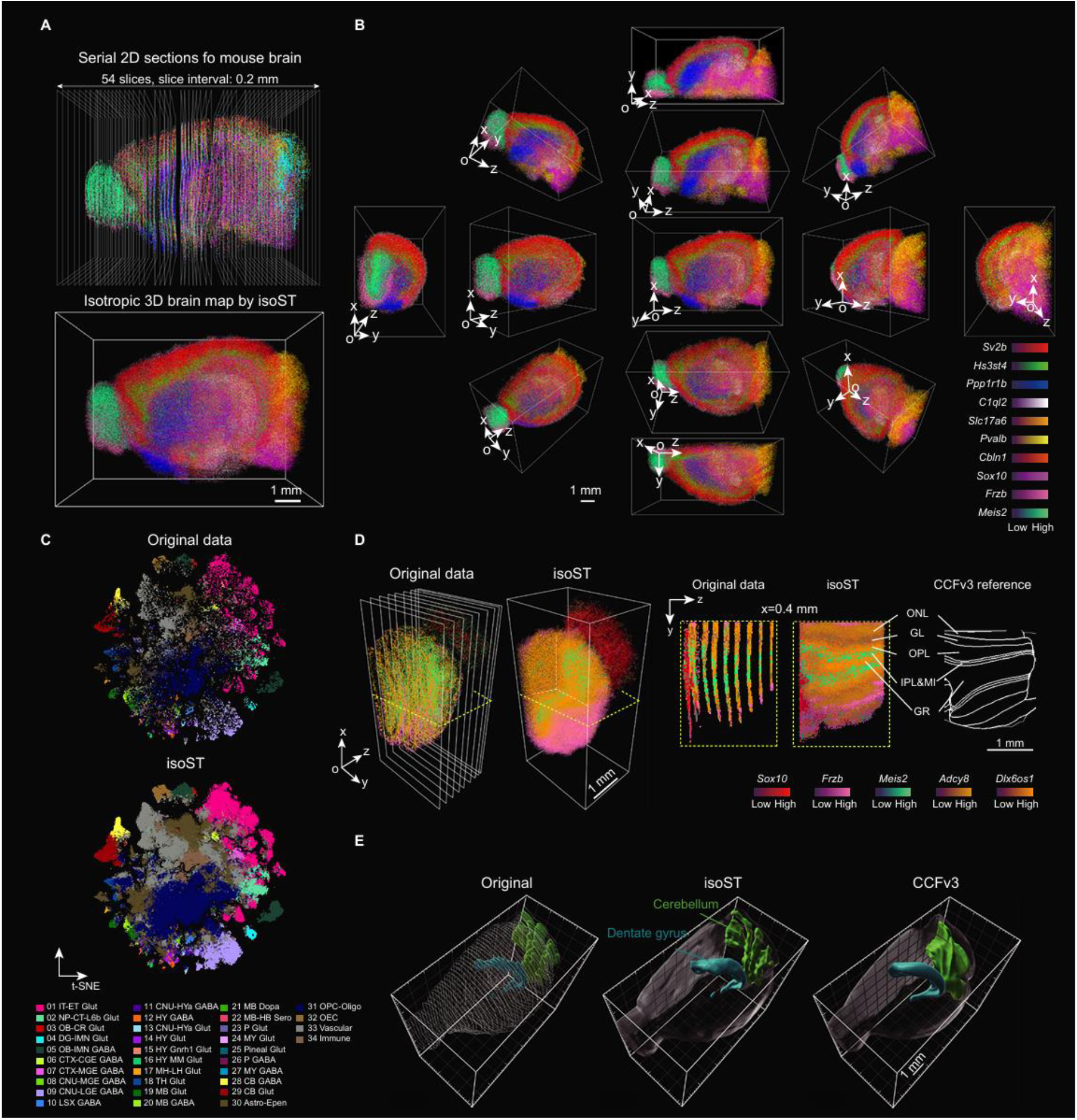
3D reconstruction of the mouse brain using isoST. (**A**) The original serial 2D sections (top) and reconstructed 3D brain map (bottom) using *n* = 54 2D mouse brain slices profiled by MERFISH. (**B**) Multi-angle views of the reconstructed 3D brain map with marker genes across major anatomical axes. (**C**) Visualization (34 subclasses) of cell transcriptional profiles in the original (top) or isoST-reconstructed (bottom) data using the same t-SNE space. (**D**) Comparison of spatial transcriptomics profiles of olfactory bulb before and after isoST reconstruction. Left: the original slices and isoST-reconstructed volume; yellow dashed boxes indicate the plane of section shown on the right. Right: transverse view at x = 0.4 mm, displaying gene expression patterns in the original slices, isoST volume, and comparing with known structures from CCFv3 reference. ONL: olfactory nerve layer, GL: glomerular layer, OPL: outer plexiform layer, MI: mitral cell layer, IPL: inner plexiform layer, and GR: granule cell layer. (**E**) Spatial visualization of dentate gyrus and cerebellum across the original annotation (left), isoST reconstruction (middle), and CCFv3 reference (right). Additional results for other regions are provided in **Supplementary Fig. 4**.

We then applied isoST to the full set of 54 slices to reconstruct a continuous, isotropic-resolution 3D transcriptomic map of the mouse brain (**Fig. 2A**, bottom). The expression of known marker genes displayed anatomically coherent and smoothly varying patterns across brain regions (**Fig. 2B**; **Movie S2**). Reconstructed slices closely matched the original profiles, preserving both broad cell type distinctions (**Fig. 2C**) and finer subclass resolution in t-SNE space(*27*) (**Supplementary Fig. 3**). To assess local structural fidelity, we compared reconstructed gene expression patterns with anatomical annotations from the Allen Mouse Brain Common Coordinate Framework version 3 (CCFv3)(*25*). In the olfactory bulb, marker genes captured the known laminar organization(*28-31*) (**Fig. 2D**, right), while reconstructed structures in the dentate gyrus, cerebellum (**Fig. 2E**), and other regions (**Supplementary Fig. 4**) aligned with well-established anatomical features. Furthermore, inspired by the BANKSY framework(*32*), we extracted 3D spatial features combining regional averages and local gradients for unsupervised clustering (**Methods**). Compared with nonspatial clustering on the isoST volume, 3D spatial clustering achieves substantially improved anatomical coherence and inter-slice consistency (**Supplementary Fig. 5**). These results demonstrate that isoST can reconstruct isotropic-resolution 3D transcriptomic landscapes from sparsely sampled 2D slices, faithfully preserving both anatomical structure and gene expression patterns.

### Validation of 3D spatial transcriptomics reconstruction across diverse organs

We next applied isoST to two additional organs: a mouse kidney and a Carnegie stage 7 (CS7) human gastrula(*23*) (**Methods**). The kidney dataset consisted of 8 serial slices profiled using Array-seq(*33*). isoST reconstructed a smooth 3D spatial continuum (**Fig. 3A**), capturing key anatomical transitions from the collecting ducts in the medulla to the bifurcating structures of the urothelium (**Fig. 3B**, top), along with additional internal features (**Fig. 3B**, bottom; **Supplementary Fig. 6**). In the CS7 gastrula dataset, comprising 82 Stereo-seq(*34*) sections, isoST generated an isotropic 3D transcriptomic map of the early human embryo (**Supplementary Fig. 7A**), enabling precise spatial localization of embryonic regions (**Fig. 3C**; **Supplementary Fig. 7B**). These results highlight the generalizability of isoST for reconstructing spatially continuous 3D gene expression and cell-type architectures across diverse organ systems.

**Figure 3.**
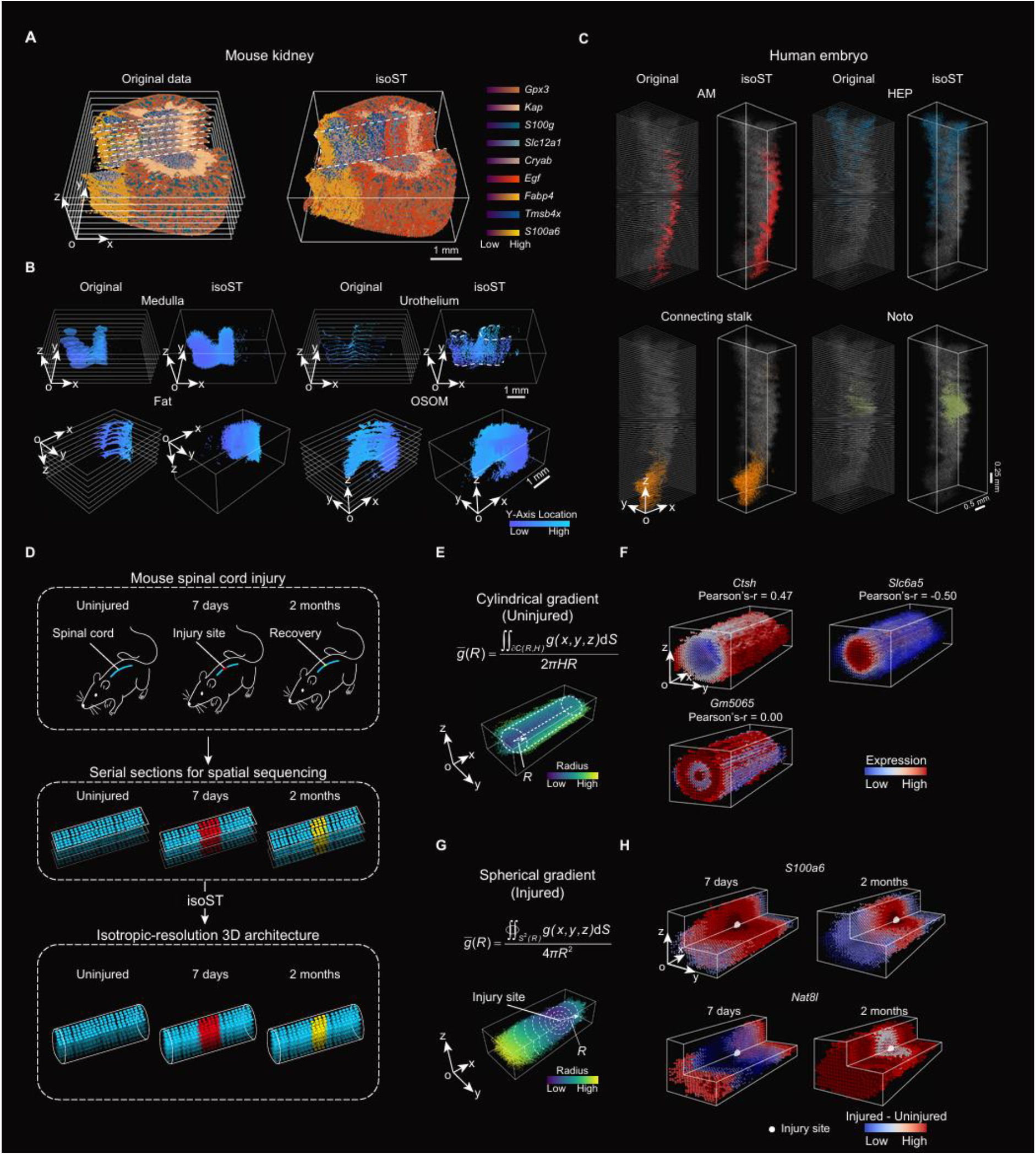
Isotropic-resolution 3D reconstruction and analysis across diverse tissues. (**A**) The original serial 2D sections (left) and reconstructed 3D map (right) for a mouse kidney using *n* = 8 2D slices profiled by Array-seq. (**B**) Visualization of original (left) or reconstructed 3D structure (right) for selected kidney regions. OSOM: outer stripe of outer medulla. (**C**) Visualization of original or isoST-reconstructed regions from a Carnegie Stage 7 (CS7) human gastrula profiled by *n* = 82 Stereo-seq sections. AM: amnion; HEP: haemato-endothelial progenitors; Noto: notochord. (**D**) Illustration of the analysis on mouse spinal cord injury dataset. Top: experimental design; middle: raw 2D slices for Visium profiling from each condition; bottom: isoST reconstruction for 3D transcriptomics profiles. (**E-F**) Cylindrical gradient analysis for the uninjured spinal cord. (**E**) Definition of cylindrical gradient of gene expressions. *R*: radius; *H*: height; *∂C*(*R, H*): the cylindrical shell. (**F**) Visualization of 3D expression patterns for example genes with positive (*Ctsh*), negative (*Slc6a5*), and uncorrelated (*Gm5065*) radial gradients. (**G-H**) Spherical gradient analysis for injured spinal cords. (**G**) Definition of spherical gradient of gene expressions. *S*^2^(*R*): a spherical surface of radius *R* centered on the injury core. (**H**) Visualization of 3D expression patterns for genes with positive (*S100a6*) or negative (*Nefl*) spherical gradient correlations at 7 days and 2 months post-injury.

### isoST reveals 3D expression gradients in spinal cord injury and recovery

The isotropic 3D transcriptomic profiles reconstructed by isoST enable unbiased analyses aligned with the intrinsic geometry of tissues and organs. To demonstrate this, we applied isoST to a mouse spinal cord injury dataset spanning three conditions: uninjured, 7 days post-injury, and 2 months post-injury(*24*) (**Fig. 3D**). Each condition included 16 Visium-profiled slices collected from four mice (**Methods**). We applied isoST to each condition and reconstructed full 3D transcriptomic volumes for analysis.

In uninjured samples, we modeled the spinal cord as a series of concentric cylindrical shells centered on the spinal canal (**Fig. 3E**) and identified genes exhibiting radial expression gradients based on their correlation with radial distance. Genes with strong positive (*Ctsh*) or negative (*Slc6a5*) correlations recapitulated known anatomical localization patterns(*24*) (**Fig. 3F**; **Supplementary Fig. 8A**; **Movie S3**).

In injured samples, we expected the impact of injury to radiate outward from the lesion site. To capture these spatial dynamics, we established a spherical coordinate system centered on the lesion (**Fig. 3G**) and identified genes showing spherical expression gradients. For example, *S100a6* was highly expressed near the lesion site, while *Nat8l* was enriched in more distal regions. Over time, the *S100a6*-positive area contracted and *Nat8l* expanded, reflecting progressive tissue repair (**Fig. 3H**; **Movie S3**).

We extended this spherical gradient analysis to all genes in the dataset, classifying them based on their correlation with distance from the lesion. Representative genes from each category illustrated diverse spatial patterns (**Supplementary Figs. 8B, C**). To gain functional insights, we performed pathway enrichment analysis on the positively and negatively correlated gene sets (**Methods**). At 7 days post-injury, upregulated pathways in distal regions were enriched for *MECP2*-regulated transcriptional programs associated with neural regeneration(*35*) (**Supplementary Fig. 9A**, left), whereas downregulated pathways near the lesion were associated with collagen formation and immune signaling (**Supplementary Fig. 9A**, right). By 2 months, distal regions exhibited signatures of restored neuronal function, while immune-related activity persisted at the lesion center (**Supplementary Fig. 9B**). Together, these results show that isoST enables geometry-aware analysis of molecular dynamics in injury and recovery across both spatial and temporal dimensions.

### isoST enables accurate reconstruction from fewer sections by integrating imaging data

Reconstructing 3D gene expression typically requires transcriptional profiling of many serial tissue sections to preserve anatomical fidelity. To reduce the number of transcriptional profiled slices without compromising reconstruction quality, we extended isoST to incorporate densely sampled histological imaging, resulting in isoST-i (**Fig. 4A**). isoST-i uses image-derived structural features to regularize the inference of intermediate layer between observed transcriptional layers, thereby constraining and enhancing reconstruction accuracy (**Fig. 4A**; **Supplementary Fig. 10**).

**Figure 4.**
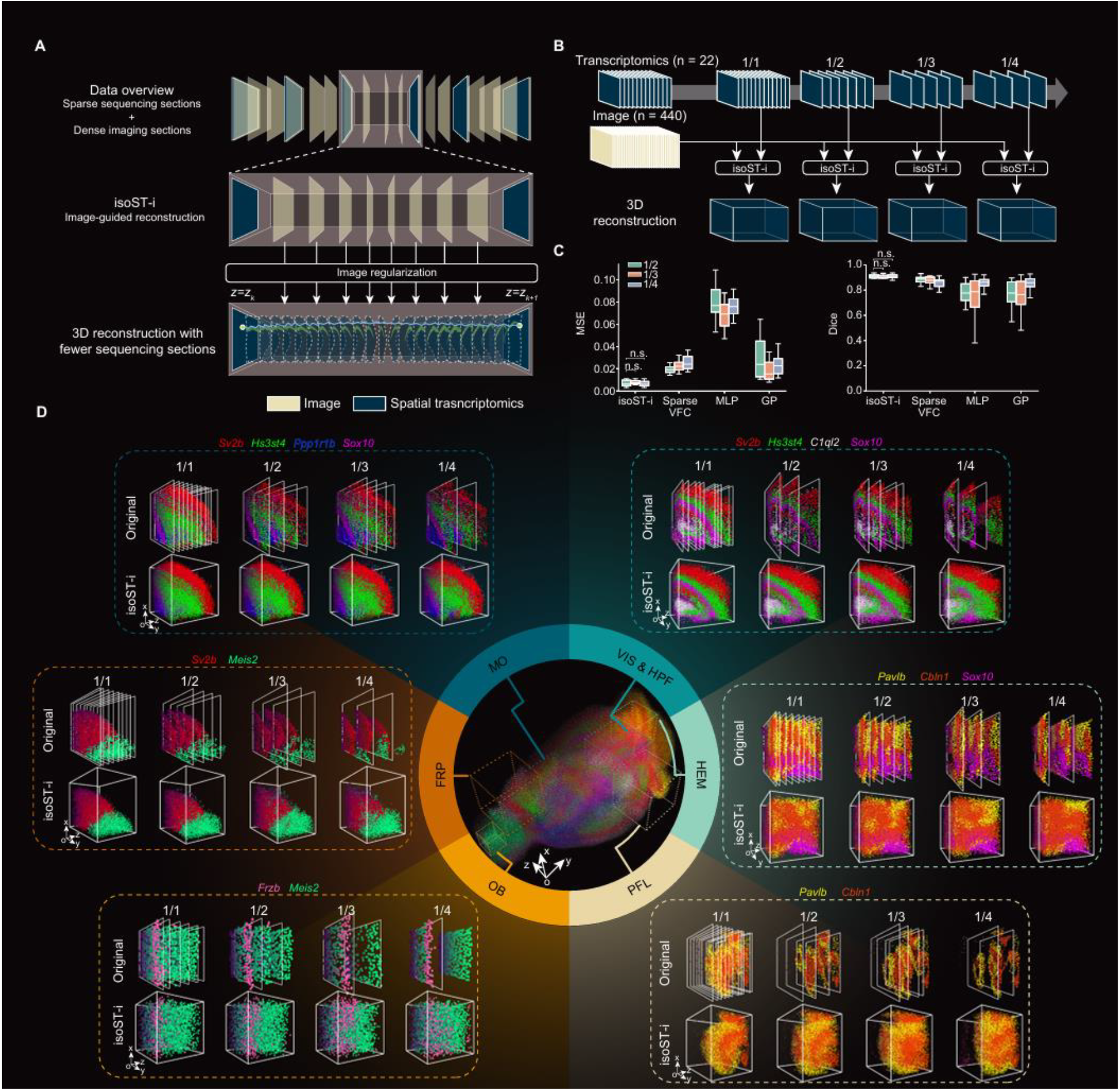
Image-guided 3D reconstruction of the mouse brain from reduced transcriptomic input. (**A**) Overview of the isoST-i framework. isoST-i combines dense image stacks to regularize the reconstruction process for accurate 3D transcriptomic reconstruction from sparsely sampled spatial omics slices. (**B**) Experimental setup for evaluating reconstruction under reduced transcriptomic input. The original dataset of 22 transcriptionally profiled tissue sections was subsampled to 1/2, 1/3, and 1/4 of the original slices while retaining the full imaging stack for guidance. (**C**) Quantitative comparison of reconstruction accuracy using mean squared error (MSE, left) and Dice coefficient (right). Box plots represent the median (center line), interquartile range (box), and full range (whiskers). Statistical significance was assessed using two-sided rank-sum tests on held-out slices; n.s.: not significant (p ≥ 0.05). (**D**) Regional gene expression analysis across six brain regions: OB (olfactory bulb), FRP (frontal pole), MO (somatomotor areas), VIS (visual areas), HPF (hippocampal formation), HEM (hemispheric domain), and PFL (paraflocculus). For each region, isoST-i reconstructions (bottom) based on varying sampling ratios (1/1, 1/2, 1/3, 1/4) are compared with the corresponding original slices (top).

As a test case, we applied isoST-i to a new MERFISH-based mouse brain dataset consisting of 22 spatial transcriptomic slices(*19*), supported by 440 histological sections from the CCFv3 atlas(*25*) (**Methods**). We progressively down sampled the transcriptomic input to 1/2, 1/3, and 1/4 of the original slices, while maintaining the same number of image slices to reconstruct isotropic 3D brain maps (**Fig. 4B Movie S4**). Across all settings, isoST-i preserved gene expression accuracy and anatomical structure, as confirmed by comparisons with held-out slices (**Fig. 4C**). Both global reconstructions (**Supplementary Fig. 11**) and detailed regional views across six brain areas (**Fig. 4D**) demonstrated that isoST-i maintained smooth boundaries and coherent architecture with only 1/3 of the original slices. At results with 1/4 of input, some regions such as the visual cortex (VIS), hippocampal formation (HPF), and frontal pole (FRP) began to show mild structural blurring.

### isoST models precise temporal continuity across whole-embryo development

In the temporal dimension, spatial transcriptomic studies face a similar challenge as in the spatial domain: only a few discrete timepoints are typically sampled due to experimental constraints, limiting temporal resolution of developmental dynamics. To address this, we extended isoST (named as isoST-t) to model temporal continuity by treating developmental timepoints as analogous to spatial “depth.” As a demonstration, we applied isoST-t to a Stereo-seq mouse embryo dataset collected at five discrete developmental stages: E10.5, E12.5, E13.5, E15.5, and E16.5(*34*) (**Methods**). isoST-t enhanced the temporal resolution to 0.1-day resolution, generating 61 gene expression slices that capture the dynamics of embryonic development (**Fig. 5A**; **Supplementary Fig. 12A**).

**Figure 5.**
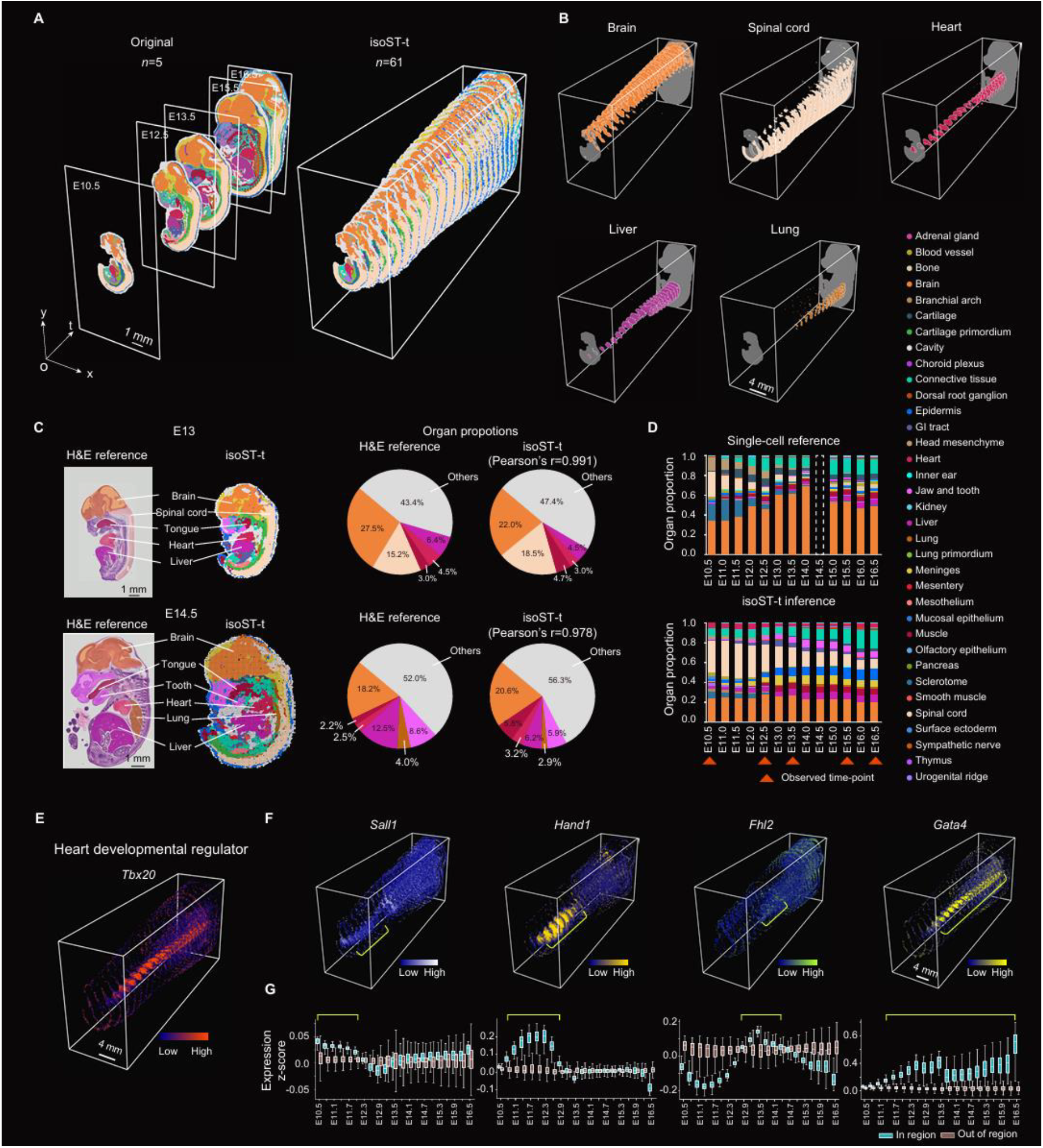
Repurpose of isoST to the dense reconstruction of spatial–temporal trajectories during mouse embryonic development. (**A**) Left: original spatial transcriptomic slices collected at five developmental timepoints (E10.5, E12.5, E13.5, E15.5, and E16.5). Right: reconstructed spatial transcriptomic profiles at 61 reconstructed timepoints using isoST-t. (**B**) Visualization of organ trajectories for five major organs (brain, spinal cord, heart, liver, and lung) across reconstructed developmental stages. (**C**) Validation of isoST-t reconstructions at unprofiled timepoints (E13 and E14.5) using histological references from the eMouse Atlas. Pie charts compare organ composition between the two sources. (**D**) Validation of isoST-t reconstructed slices by comparing against a single-cell RNA-seq atlas spanning 12 timepoints. Stacked bar plots show organ composition inferred from isoST-t versus the single-cell reference. Timepoints without single-cell data are indicated by dashed white boxes. (**E-F**) Visualization of expression patterns for genes that were consistently expressed during development (**E**) or dynamically regulated during across developmental stages for multiple organ regions. Yellow bars indicate peak expression windows. Blue and gray denote whether cells are located within or outside the corresponding organ regions.

To assess the accuracy of the inferred intermediate stages, we compared the reconstructed embryo sections at unprofiled timepoints (E13 and E14.5) to real hematoxylin and eosin (H&E) histological sections(*36*). The reconstructed spatial structures showed high consistency (r=0.99 for E13 and 0.97 for E14.5) with organ-level annotations across multiple tissues, including the brain, spinal cord, tongue, heart, liver, and lungs (**Fig. 5C**). We then evaluated how well the reconstructed organ composition aligned with a single-cell transcriptomic atlas of mouse embryogenesis that spans 12 developmental stages(*37*). Despite differences in experimental platforms and sampling strategies, the relative proportions of major organs in isoST-t reconstructions closely matched those derived from the single-cell data, with high Pearson correlation across time points (>0.6 for all timepoints; **Fig. 5D**; **Supplementary Fig. 12B**).

The temporally dense reconstructions enabled fine-grained analysis of gene expression dynamics. We first identified organ-specific genes showing consistent differential expression across stages (**Methods**), including known developmental regulators such as *Tbx20* (heart)(*38, 39*), *Kat2a* (brain)(*40*), and *Hoxb7* (spinal cord)(*41*) (**Fig. 5E**; **Supplementary Fig. 12C**). We also explored stage-specific regulatory shifts for organs (**Methods**). For instance, *Sall1* was enriched in early-stage hearts but decreased after E13.5, consistent with its role in cardiac progenitor proliferation and compact myocardium formation(*42, 43*) (**Fig. 5G**, first panel). *Hand1* peaked at E12.0, marking a transitional phase in chamber development(*44, 45*) (**Fig. 5G**, second panel). *Fhl2* was broadly repressed except for a sharp rise at E14.5, aligning with its role in myocardial structural remodeling(*46-48*) (**Fig. 5G**, third panel). *Gata4* steadily increased across stages, peaking at E16.5, consistent with its function in cardiac maturation and valve development(*49-51*) (**Fig. 5G**, fourth panel). These results demonstrate that isoST-t enables high-resolution tracking of gene regulation dynamics across embryonic time.

## Discussion

Capturing the inherent three-dimensional organization of gene expression within intact tissues has long been a central goal in molecular and developmental biology. While recent spatial transcriptomic technologies have made significant advances, they remain constrained by practical limitations, such as the need for dense tissue sectioning, high sequencing costs, and incomplete spatial coverage. In this study, we present isoST, a practical and generalizable computational framework that enables isotropic-resolution 3D reconstruction of gene expression landscapes from sparsely sampled 2D spatial omics data. We show that isoST can faithfully recover known anatomical and transcriptional structures across multiple organs through quantitative comparison with held-out slices and anatomical references. Extensions of isoST enable integration with dense imaging data to reduce the number of required transcriptomic slices, lowering experimental burden while preserving reconstruction accuracy. Additionally, isoST can model temporal continuity across developmental stages, providing fine-grained resolution of gene expression dynamics over time.

While isoST performs robustly across various datasets, its accuracy depends on several factors, including tissue complexity, the number of spatial transcriptomic slices, and data quality. The method is well suited for interpolating continuous transitions between known spatial or temporal states, but it does not predict entirely novel structures that fall outside the sampled input. Therefore, isoST complements—but does not replace—experimental strategies designed to capture rare or previously unobserved features.

A natural next step is to improve the adaptability of isoST to different biological contexts. An important challenge is to determine how many sections are required to ensure reliable reconstruction and to quantify the extent to which structural information from imaging can compensate for sparse molecular data. Moreover, extending the model to jointly capture both spatial and temporal dynamics in a unified 4D framework would enable the reconstruction of tissue morphogenesis with high resolution across both space and time.

Together, isoST provides a practical solution for reconstructing high-resolution 3D transcriptomic maps from sparse data. By reducing reliance on dense profiling and enabling integration with auxiliary information, it serves as a flexible tool for spatial biology and large-scale mapping efforts such as the Human Cell Atlas.

## Materials and methods

### A stochastic differential equation model for isotropic 3D spatial transcriptomics

In a three-dimensional (3D) spatially resolved transcriptomics dataset, one tissue is sectioned into *K* serial slices for sequencing. We define the (*x, y*) as directions for each individual slice and the *z* as the tissue depth, along which multiple slices are arranged. For a two-dimensional (2D) slice at the depth *z*_*k*_, the spatial sequencing profiles are represented as a function of spatial positions 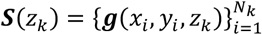, where *N*_k_ is the number of cells (or spots) in this tissue slice, and ***g***(*x*_*i*_, *y*_*i*_, *z*_*k*_) ∈ ℝ^*G*^ is the gene expression at the position (*x*_*i*_, *y*_*i*_, *z*_*k*_). The goal is to infer the spatial expression ***g***(*x, y, z*^′^) for unprofiled tissue depths *z*^′^ ∈ [*z*_1_, *z*_*K*_], thereby construct a full 3D transcriptional continuum for the tissue.

However, compared to the (*x, y*) directions, where thousands to millions of spatial positions are densely measured in a dataset, the *z*-direction usually only includes a sparse set of depth values. This imbalanced spatial information makes it challenging to directly regress or interpolate new tissue layers.

Here, we leverage the tissue’s inherent spatial continuity and model the tissue slice at an unprofiled depth *z*_*k*_ + Δ*z* as a gradual extension from an existing, nearby slice at the depth *z*_*k*_. To achieve this, we make use of two stochastic differential equations (SDEs) to model the tissue shape and expression in the 3D space, respectively:

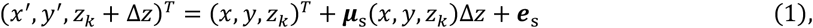

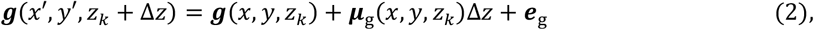

where the (*x, y, z*_*k*_) and (*x*^′^, *y*^′^, *z*_*k*_ + Δ*z*) are measured (at layer *z*_*k*_) and predicted (at layer *z*_*k*_ + Δ*z*) cell/spot positions, while ***g***(*x, y, z*_*k*_) and ***g***(*x*′, *y*′, *z*_*k*_ + Δ*z*) are their corresponding expression profiles; the drift coefficient ***μ***_s_(⋅) ∈ ℝ^3^ and ***μ***_g_(⋅) ∈ ℝ^*G*^ are learnable gradient functions modeling the spatial change of the gene expressions and tissue shapes along with the tissue depth, respectively; the diffusion terms ***e***_s_ ∈ ℝ^3^ and ***e***_g_ ∈ ℝ^*G*^ are white Gaussian noise.

With a small step Δ*z*, the first SDE models how the cell distribution in the (*x, y*)-plane changes along with depth *z*_*k*_ while the second SDE models the corresponding expression change. With the depth accumulation of multiple steps, the SDE model incrementally constructs (near-)continuous expression transitions along the *z*-axis between two profiled layers, thus establishing a spatial continuum of transcriptomics.

To learn the gradient functions, we employ graph neural networks (GNNs)(*52, 53*) *f*_s_ and *f*_g_ to aggregrate the neighboring profiles around (*x, y, z*_*k*_) to predict how tissue shape and gene expression change as the tissue depth shifts by Δ*z*:

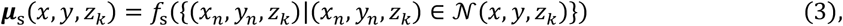

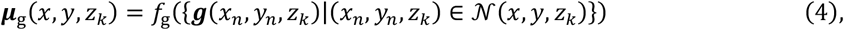

where 𝒩(*x, y, z*_*k*_) is all profiled cell/spots spatially around the location (*x, y, z*_*k*_) from slice at depth *z*_*k*_.

### Model optimization

Starting from a profiled tissue layer at *z*_*k*_, we integrate **Eqs. 1** to **4** to infer the tissue profile at the layer *z*_*k*+1_:

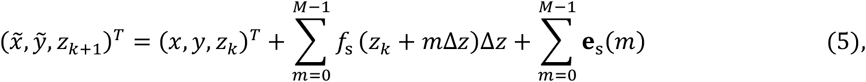

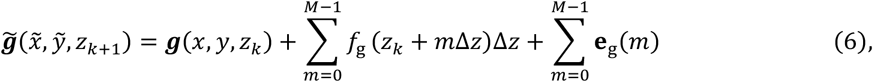

where 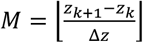 is the number of steps used from depth *z*_*k*_ to 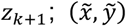 is the inferred spatial distribution of cells at the depth *z*_*k*+1_ and 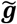 is the inferred gene expressions. Therefore, we can formulate the loss function by aligning both the shape and expression levels between predicted and profiled data. Specifically, we use the Wasserstein distance to measure the cell position distribution difference while using the mean squared error (MSE) for the accuracy of expression profiles. The loss at depth *z*_*k*+1_ is defined as:

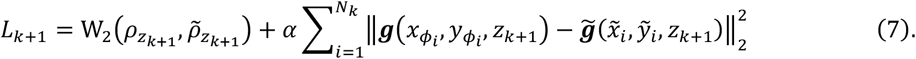

Here, the shape alignment term W_2_(⋅) represents the Wasserstein distance between the observed and predicted cell positions; 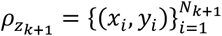 and 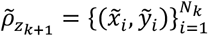 are profiled or inferred cell position distributions. In the expression alignment term, 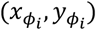 is the nearest cell position in the real data to the position 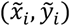 in the inferenced data. The hyperparameter *α* balances the contributions of two terms and is determined empirically from experiments (**Supplementary Table 1**). The total loss function is the summation over all layers. In the implementation, we also consider the backward inference from layer *z*_*K*_ towards *z*_1_ and the final loss function is formulated as:

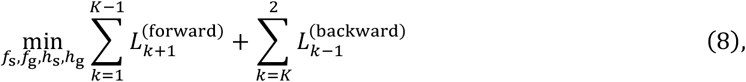

where 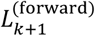 is the rewritten form of *L* _*k*+1_in **Eq. 7**. The backward functions and corresponding GNNs (*h*_*s*_ and *h*_*g*_) are defined accordingly (**Supplementary Note 1**).

The multi-step iterative process in **Eqs**. 5 and 6 is implemented using a neural ODE framework(*54*) and optimized with the NAdam optimizer(*55*) at a learning rate of 0.01. We employ a three-step strategy to ensure both accuracy and efficiency: (1) Minimize the shape alignment term to accurately predict the spatial coordinates of the slices; (2) Minimize the expression alignment term to closely match the actual transcriptomic profiles; (3) Minimize the overall loss function to achieve comprehensive slice characterization.

### Reconstruction of 3D spatial transcriptomics profiles

After optimization, isoST infers the tissue profile at any intermediate depth *z* ∈ (*z*_*k*_, *z*_*k*+1_) by integrating the learned drift functions from the nearest observed slices, as defined in **Eqs. 5**-**6**. Specifically, the forward integration propagates from *z*_*k*_ to *z*, while the backward integration proceeds from *z*_*k*+1_ to *z*, using symmetric drift modules *h*_*s*_ and *h*_*g*_. The number of integration steps is determined by ⌊(*z* − *z*_*k*_)/Δ*z*⌋ for the forward path, and analogously for the backward path. The final reconstruction at depth *z* is defined as the union of both paths:

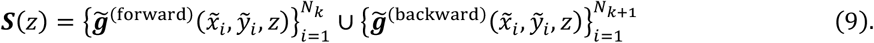

To generate a unified 3D representation, we aggregate all reconstructed slices into a unified tissue-level set: ***S*** = ⋃_*z*∈***Z***_ ***S***(*z*), where *Z* = {*z*_1_, *z*_1_ + Δ*z*, …, *z*_*K*_} is the set of all observed and reconstructed depths. To construct a volumetric gene expression tensor ***V***, we define a regular voxel grid with spacing Δ*v* in each dimension. For each voxel at position (*x, y, z*), the expression vector is computed as the average value of all expressions located within:

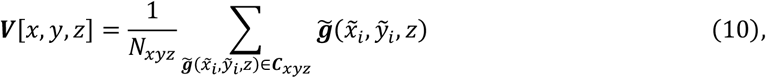

where ***C***_*xyz*_ ⊂ ***S*** denotes the set of observed and reconstructed data points falling inside the cube centered at (*x, y, z*) with side length Δ*v*, and *N*_*xyz*_ = |***C***_*xyz*_|.

### Image-guided isoST reconstruction

We extended isoST to incorporate image-derived structural priors. For depths 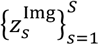 with available images, we introduce two regularization terms to guide prediction of spatial coordinates and gene expression at intermediate slices.

For spatial regularization, we convert image data into point clouds based on estimated cell densities, forming reference shapes 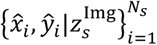. For expression regularization, we align imputed profiles with image-derived feature maps ***Î***_*s*_ via a linear projection ***W*** trained on paired image-transcriptome data. The image-based loss is:

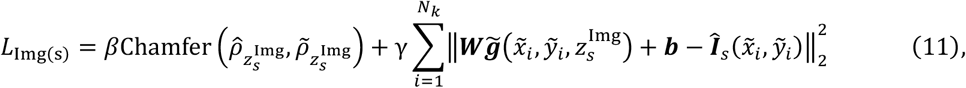

where Chamfer(·,·) measures distance between cell coordinate distributions from image 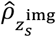 and imputed slice 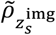 (*56*); the projection matrix ***W*** ∈ ℝ^*F*×*G*^ and bias ***b*** ∈ ℝ^*G*^ are obtained from a pretrained linear regression model based on paired image-expression data; *β* and *γ* balance the two terms (**Supplementary Table 2**). The total loss integrates image regularization with forward and backward inference:

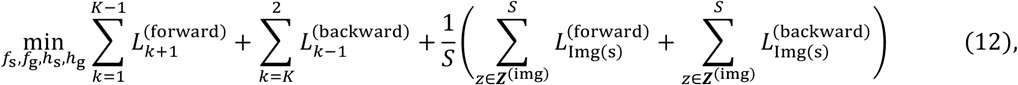

where 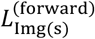 denotes the image regularization term in the forward direction (as defined in **Eq. 11**), while 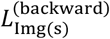 is its counterpart in the backward pass.

## Datasets

### MERFISH mouse brain dataset

The mouse brain was obtained from a male mouse aged approximately 56–62 days (sample ABCA-2 from the original dataset). The brain was sectioned into 54 coronal slices at intervals of 0.2 mm, and each slice was individually profiled using the MERFISH platform. In total, the expression of 1,122 genes was measured across 1,056,520 cells. Gene expression counts were normalized to the total counts per cell, log1p-transformed, and projected onto the top 50 principal components (PCs). Marker genes for each anatomical region were identified using the *rank_genes_groups* function from Scanpy (v1.10.3). Cell annotations were obtained from original publication.

### Array-seq mouse kidney dataset

The mouse kidney was obtained from a mouse aged approximately 6–8 weeks and sectioned into eight slices at 0.1 mm intervals. Transcripts were directly profiled across all sections using the Array-seq platform. In total, 15,144 genes were measured across 92,769 spatial spots. Normalized expression data from the original publication were used and the 3,000 most highly variable genes were selected for analysis. Marker genes for each region were identified as previously described. Tissue annotations were obtained from the authors’ analysis.

### Stereo-seq human embryo dataset

The sample was collected from a male human embryo corresponding to gestational week 5. The embryo was sectioned into 82 slices at 0.01 mm intervals and profiled using the Stereo-seq platform. In total, 25,833 genes were measured across 28,804 spatial spots. Genes detected in fewer than 10 spots were excluded, retaining 22,079 genes for downstream analysis. Gene expression values were normalized to a total count of 10,000 per spot and log1p-transformed. The 2,000 most highly variable genes were selected and projected onto the top 50 PCs. Marker genes for each region were identified as previously described. Tissue annotations were obtained from a previous publication.

### Visium mouse spinal cord dataset after injury

The mouse spinal cord dataset was collected from mice representing three injury states: uninjured, 7 days post-injury, and 2 months post-injury following spinal cord crush. For each state, four mice were used, and four spinal cord sections were collected per mouse, yielding 16 sections per state. In total, 26,136 genes were measured across 37,558 spatial spots using the Visium platform. Gene expression values were normalized to the median total count per spot, log1p-transformed, and projected onto the top 50 PCs. The 2,000 most highly variable genes were selected. Tissue domains were identified using the Leiden clustering algorithm (resolution = 0.2), and marker genes were identified as previously described.

### Image-guided reconstruction

The mouse brain was obtained from a male mouse aged approximately 56–62 days (sample ABCA-3 from the original dataset). The brain was sectioned into 22 coronal slices at intervals of 0.2 mm, and each slice was individually profiled using the MERFISH platform.

The image data were obtained from the Allen Mouse Brain Common Coordinate Framework version 3 (CCFv3). We extracted coronal sections aligned with the orientation of the MERFISH brain dataset. To derive image features, we computed the mean and standard deviation of voxel intensities using kernels of sizes 3, 5, 7, and 9, along with the raw intensity values, resulting in a 10-dimensional feature map. For shape regularization, we retained voxels with intensity values greater than 6 and sampled voxels with probabilities proportional to their intensity to construct a shape prior.

### Stereo-seq mouse embryo dataset for temporal modeling

The mouse embryo dataset included five developmental stages: E10.5, E12.5, E13.5, E15.5, and E16.5. In total, 392,705 spatial spots were profiled using the Stereo-seq platform. In the original publication, 844 gene regulons were defined for developmental analysis. To correct batch effects across sections, the data were normalized using the *combat* function(*57*) from Scanpy (v1.10.3) and projected onto the top 50 PCs. Organ-enriched genes were identified using the *rank_genes_groups* function across stages, and stage-specific genes were obtained by applying the same function to each slice individually. In both cases, the top 1 differentially expressed genes were selected for downstream analysis.

### H&E image embryo dataset for reconstruction evaluation

Hematoxylin and eosin (H&E) stained reference sections were obtained from the eMouseAtlas database(*36*). Samples from embryonic stages E13 (montage_29aa) and E14.5 (montage_33ba) were used to evaluate the inferred structures at the same timepoints. Organ regions were manually annotated based on morphological features by visual inspection.

### scRNA-seq mouse embryo dataset for reconstruction evaluation

Mouse embryo single-cell RNA sequencing (scRNA-seq) data spanning stages E10.5 to E16.5 at 0.25-day intervals were used for reconstruction evaluation. A total of 188 cell types were annotated in the original dataset. Major cell types corresponding to specific organs (**Supplementary Table 1**) were used to estimate organ-level cell proportions across developmental stages.

## Experiment setup

### Cell type identification

A K-nearest neighbors (KNN) classifier was trained using existing cell type annotations from each dataset and applied to predict cell types in the reconstructed 3D transcriptomic data. For the mouse brain dataset, predicted labels were further refined using the *LabelPropagation* algorithm (v2.5.2) for visualization.

### 3D spatial feature extraction from isoST-reconstructed volume

We designed 3D spatial features to capture both regional expression trends and local gradients. To obtain regional averages, we first smoothed the isoST-reconstructed volume (represented by the top 50 principal components) using a 3D Gaussian kernel with standard deviation σ = (1.5, 1.5, 1.5), implemented via the *gaussian_filter function* from scipy.ndimage (v1.12.0). To extract the local spatial gradients, we next computed directional gradients by applying the Sobel operator(*58*) along the *x, y*, and *z* axes using the *convolve* function on the smoothed volume from scipy.ndimage (v1.12.0). The resulting gradient features were concatenated with the smoothed expression values and the 3D spatial coordinates, yielding a 253-dimensional feature vector per voxel. Finally, the full feature volume was projected onto 10 principal components for subsequent clustering.

### Unsupervised clustering on isoST-reconstructed volume

We performed unsupervised clustering on both the isoST-reconstructed volume and the extracted 3D spatial features using the Leiden algorithm(*59*) with a resolution parameter of 1.0 and 2 iterations. Clustering was implemented using the *leiden* function from the Scanpy package (v1.10.3).

### Spatial gradient analysis

To quantify spatial expression gradients in the spinal cord, we modeled tissues as concentric shells. For uninjured samples, cylindrical shells were centered on the spinal canal to assess radial gradients. In injured samples, to isolate injury-induced spatial changes from the baseline expression in healthy tissue, we subtracted the gene expression profiles of uninjured tissue from the corresponding regions in injured tissue. Subsequently, spherical shells were centered on the lesion core to capture outward inflammatory spread. Within each shell, gene expression was averaged and correlated with radial distance using Pearson’s correlation. To interpret spatially regulated genes, pathway enrichment analysis was performed on the top 15 genes with the strongest positive and negative correlations in each condition.

### Organ annotation for mouse embryo dataset

For the mouse embryo dataset, organ regions from reconstructed tissue slices were inferred based on existing annotations in previous publication(*34*). Labels from the profiled slices were propagated to adjacent, unprofiled sections using spatial links established by the shape gradient function in isoST-t. The resulting annotations were then discretized onto a 256 × 256 spatial grid and cell type labels were assigned by majority vote among the spots falling within that spatial bin.

## Comparing methods

All methods were benchmarked using the same input features and ran on a Linux server running Ubuntu 20.04.3 LTS with an Intel Xeon(R) 6226R CPU and an NVIDIA GeForce RTX 3090 GPU (Driver Version: 470.63.01, CUDA Version: 11.4).

### Sparse Vector Field Consensus (Sparse VFC)(*26*)

interpolates latent spatial transcriptomic slices by aligning profiles from adjacent tissue sections using a sparse vector field model. The method was implemented using the *SparseVFC* package in R (v4.2.3). The following hyperparameters were used: number of control points = 16, maximum iterations = 500, energy change rate threshold = 0.9, kernel parameter = 0.1, regularization parameter = 3, matching probability threshold = 0.75, matching probability weight = 10, energy convergence threshold = 1e−5, and minimum probability = 1e−5.

### Gaussian Process (GP)

models gene expression as a continuous, non-parametric function of spatial coordinates, providing both predictions and uncertainty estimates. We used the Sparse Variational Gaussian Process (SVGP)(*60*) approach for computational scalability, implemented with the ApproximateGP module from the *gpytorch* Python package (v1.13). For all datasets, we used the same parameter settings: 20 training iterations, batch size = 1024, number of inducing points = 512, optimizer = Adam, learning rate = 0.01, and variational ELBO as the loss function.

### Multilayer Perceptron (MLP)

models nonlinear mappings from spatial coordinates to gene expression using a feedforward neural network. The method was implemented in *PyTorch* (v1.12.0). For each spatial omics dataset, we used the same configuration: maximum iterations = 1000, batch size = 2000, learning rate = 1e−4, three hidden layers with 256 units each, optimizer = Adam, and weighted mean squared error as the loss function.

isoST is implemented in Python with PyTorch (v.1.12.0), PyG (v2.5.2), GeomLoss (v0.2.6), and torchdiffeq (v0.2.4). We provided detailed instructions and a demonstration to run the model on GitHub (details were provided at https://github.com/deng-ai-lab/isoST).

## Evaluations

### Expression accuracy evaluation with mean square error (MSE)

The mean squared error (MSE) quantifies the average squared difference between predicted and reference vectors under the l_2_ norm. Before evaluation, both vectors were min–max normalized.

MSE was computed using the mean_squared_error function from the *sklearn*.*metrics* module (v1.2.0).

### Shape accuracy evaluation with Dice coefficient

The Dice coefficient quantifies spatial overlap between predicted and ground truth regions. It is defined as twice the intersection area divided by the sum of the predicted and reference areas. A Dice score approaching 1 indicates strong agreement in spatial shape. It was computed as *Dice* = 2|*A* ∩ *B*|/(|*A*| + |*B*|), where *A* and *B* are the predicted and reference spatial coordinates.

### Organ proportion evaluation with Pearson’s correlations

To evaluate whether isoST-generated slices accurately captured organ composition, we computed Pearson’s correlation coefficients between predicted and reference organ proportions at each developmental stage. Correlations were computed using the *pearsonr* function from *scipy*.*stats* (v1.12.0).

## Software used in this study

The 3D volumes of the mouse brain were rendered by Imaris (Bitplane, v10.0.0) software. The 3D volumes of the mouse spinal cords were rendered by vedo Python package (v2025.5.3) and Plotly Python package (v5.24.1).

Software packages used in the study can be accessed via following links: Sparse VFC: https:/rdrr.io/cran/SparseVFC/; Gaussian Process: https://gpytorch.ai/; Multilayer Perceptron: https://pytorch.org/.

## Data availability

### Spatial omics datasets

All datasets used in this study are publicly available from the following sources. The mouse brain dataset (processed MERFISH data) is available from the Allen Brain Institute: https://alleninstitute.github.io/abc_atlas_access/descriptions/Zhuang_dataset.html. The mouse kidney dataset (Array-seq) is available in the Gene Expression Omnibus (GEO) under accession number GSE266244. The human embryo dataset (Stereo-seq) can be accessed at https://cs7.3dembryo.com/#/download. The mouse spinal cord dataset (Visium) is available in GEO under accession number GSE234774. The mouse embryo dataset (Stereo-seq) is available from the MOSTA database: https://db.cngb.org/stomics/mosta/.

### Reference datasets for evaluation

The CCFv3 dataset (reference image data) can be accessed via the Allen Brain Reference Atlases portal: https://portal.brain-map.org/. The H&E embryo image dataset used for reconstruction evaluation is available from eMouseAtlas. The sample from embryonic stage E13 can be accessed at https://www.emouseatlas.org/emap/eHistology/kaufman/index.php?plate=29&image=29aa, and the E14.5 sample at https://www.emouseatlas.org/emap/eHistology/kaufman/index.php?plate=33&image=33ba. The mouse embryo single-cell RNA-seq dataset used for evaluation is available in GEO under accession number GSE228590.

## Supplementary Materials

**Supplementary Note 1 | Backward progression of isoST.**

**Supplementary Table 1 | Cell Type To Organ Mapping**

**Supplementary Figure 1.**
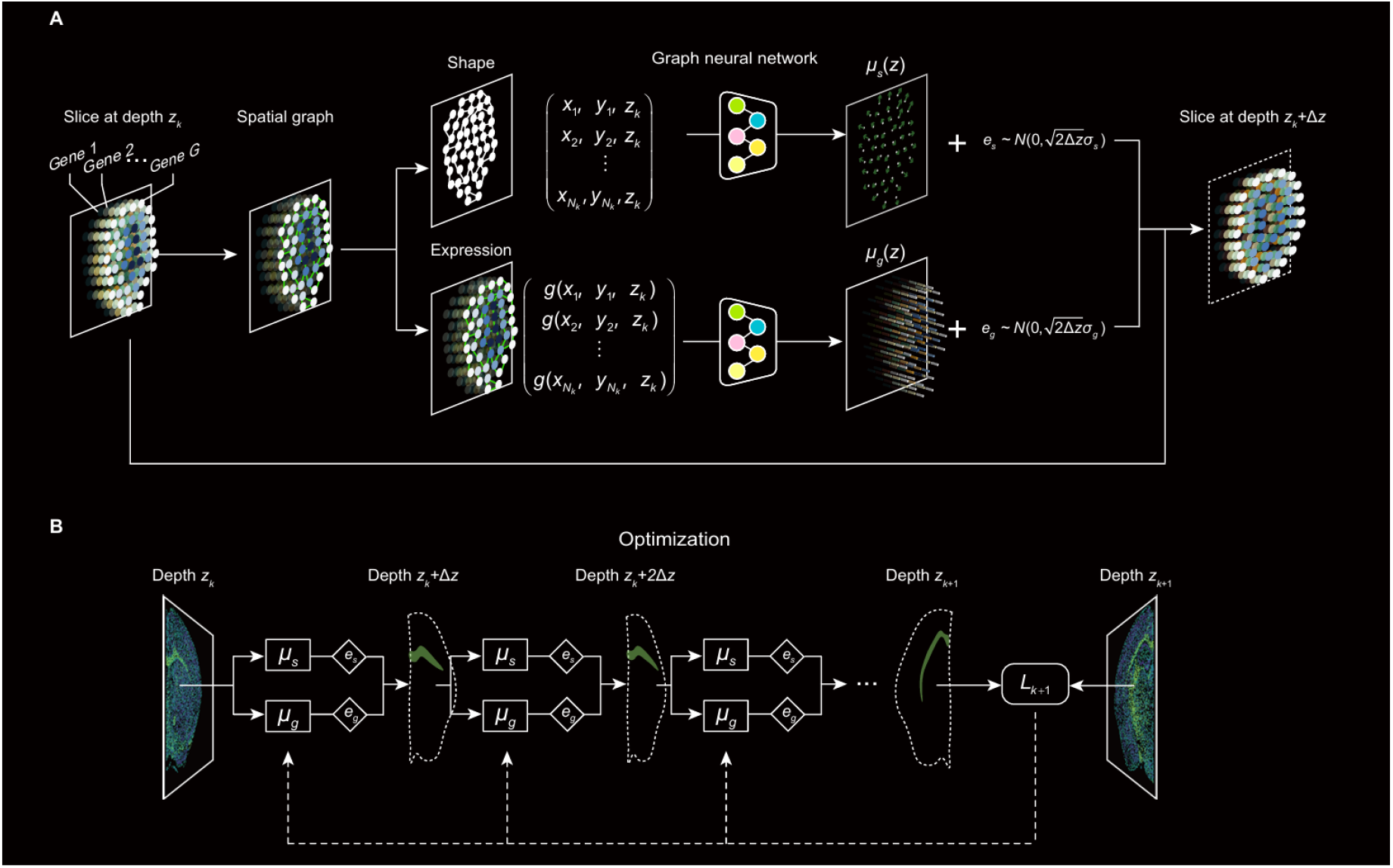
isoST structure and optimization. (**A**) Illustration of the isoST inference process from a profiled slice at depth ***z***_***k***_ to the next layer ***z***_***k***_ + **Δ*z***. A spatial graph is constructed using data point coordinates from the input slice. Two graph neural networks are then applied to predict the shape gradient ***μ***_**s**_(***z***) and the expression gradient ***μ***_**g**_(***z***). These gradients are used to generate the reconstructed slice at depth ***z***_***k***_ + **Δ*z***. (**B**) Inference between two observed slices. Starting from the slice at ***z***_***k***_, isoST iteratively reconstructs intermediate layers by applying the shape and expression inference modules ***μ***_**s**_ and ***μ***_**g**_ over small increments **Δ*z***. The reconstructed expression at the next observed slice ***z***_***k***+**1**_ is compared to the measured profile to compute the loss.

**Supplementary Figure 2.**
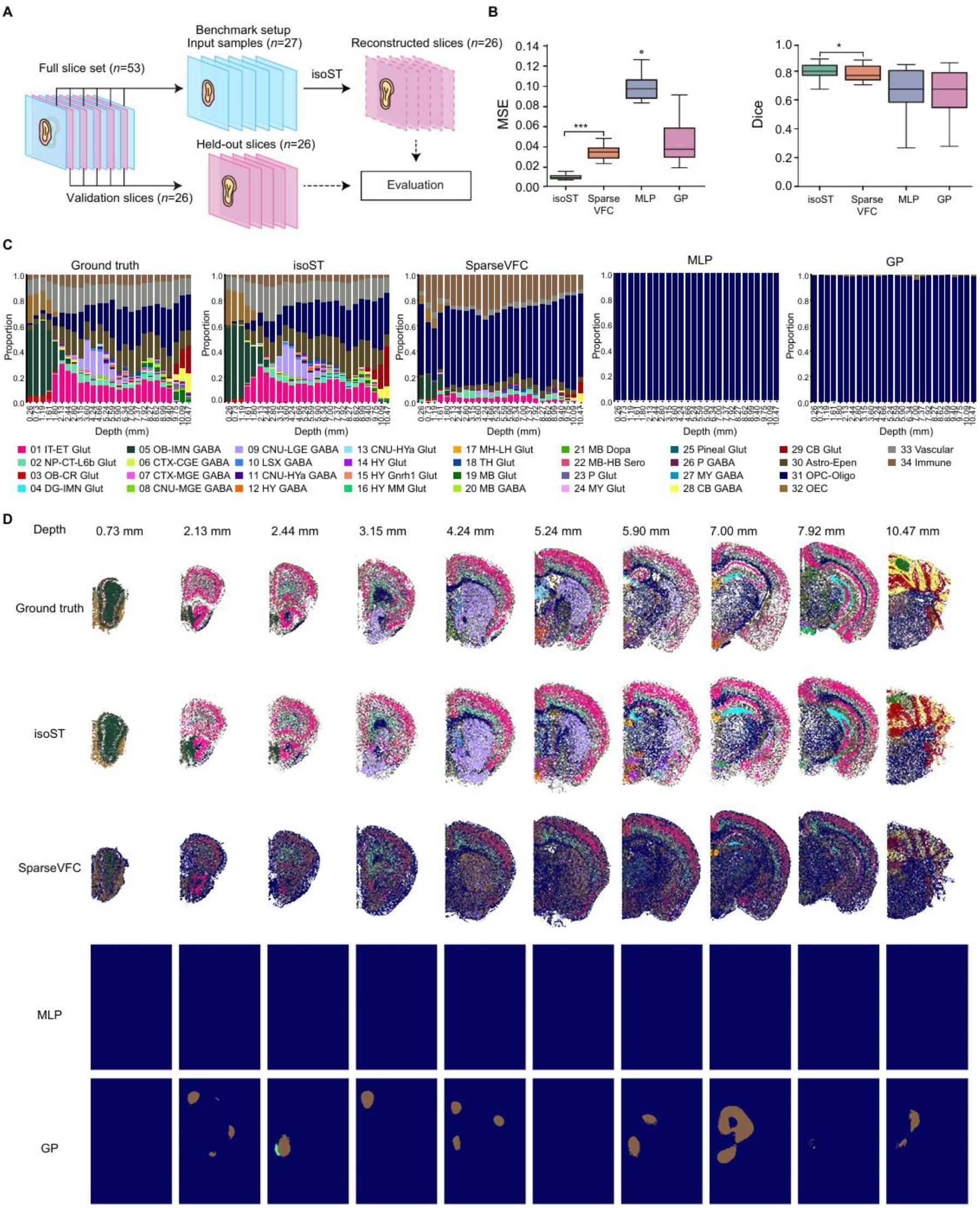
Performance evaluation on the MERFISH mouse brain dataset. (**A**) Benchmark setup. Of the 53 total slices, 27 were selected at alternating positions as input, while the remaining 26 interleaved slices were held out for evaluation. (**B**) Quantitative assessment using mean squared error (MSE) and Dice coefficient. Box plots show the median (center line), interquartile range (box), outliers (open circles), and full range (whiskers), computed across the 26 imputed slices. Statistical significance was determined using a two-sided rank-sum test: *: 1E−2 ≤ p < 0.05; ***: p < 1E−3. (**C**) Cell type composition along the depth axis, comparing the ground truth with predictions from each method. (**D**) Spatial distribution of cell types at representative depths across the brain.

**Supplementary Figure 3.**
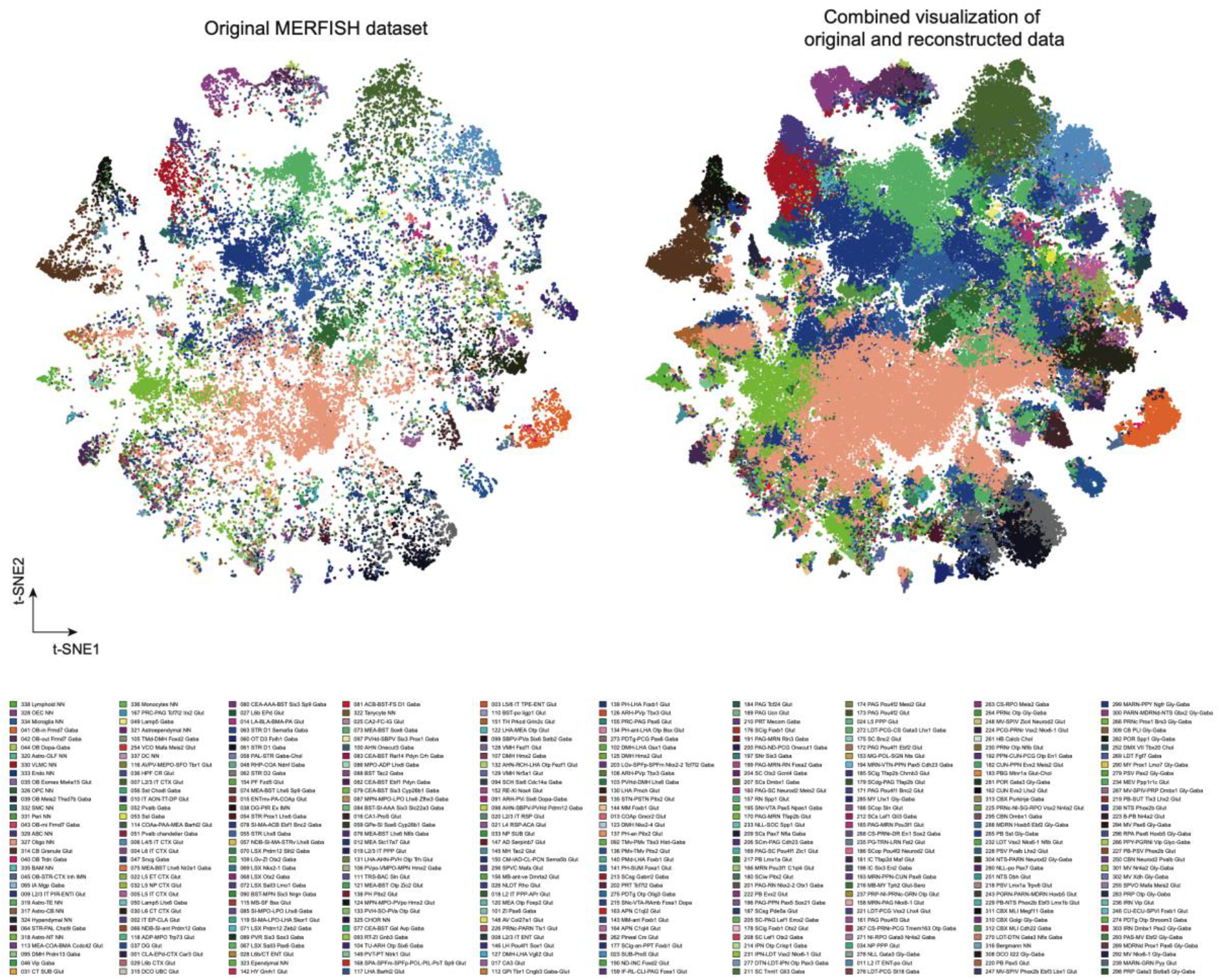
Subclass-level comparison of cell embeddings before and after isoST imputation. t-SNE visualization of 338 cell subclasses from the MERFISH mouse brain dataset. Left: embeddings computed from the original transcriptomic slices. Right: embeddings derived from isoST-reconstructed slices. Each point represents a single cell and is colored according to its annotated subclass identity (**Methods**).

**Supplementary Figure 4.**
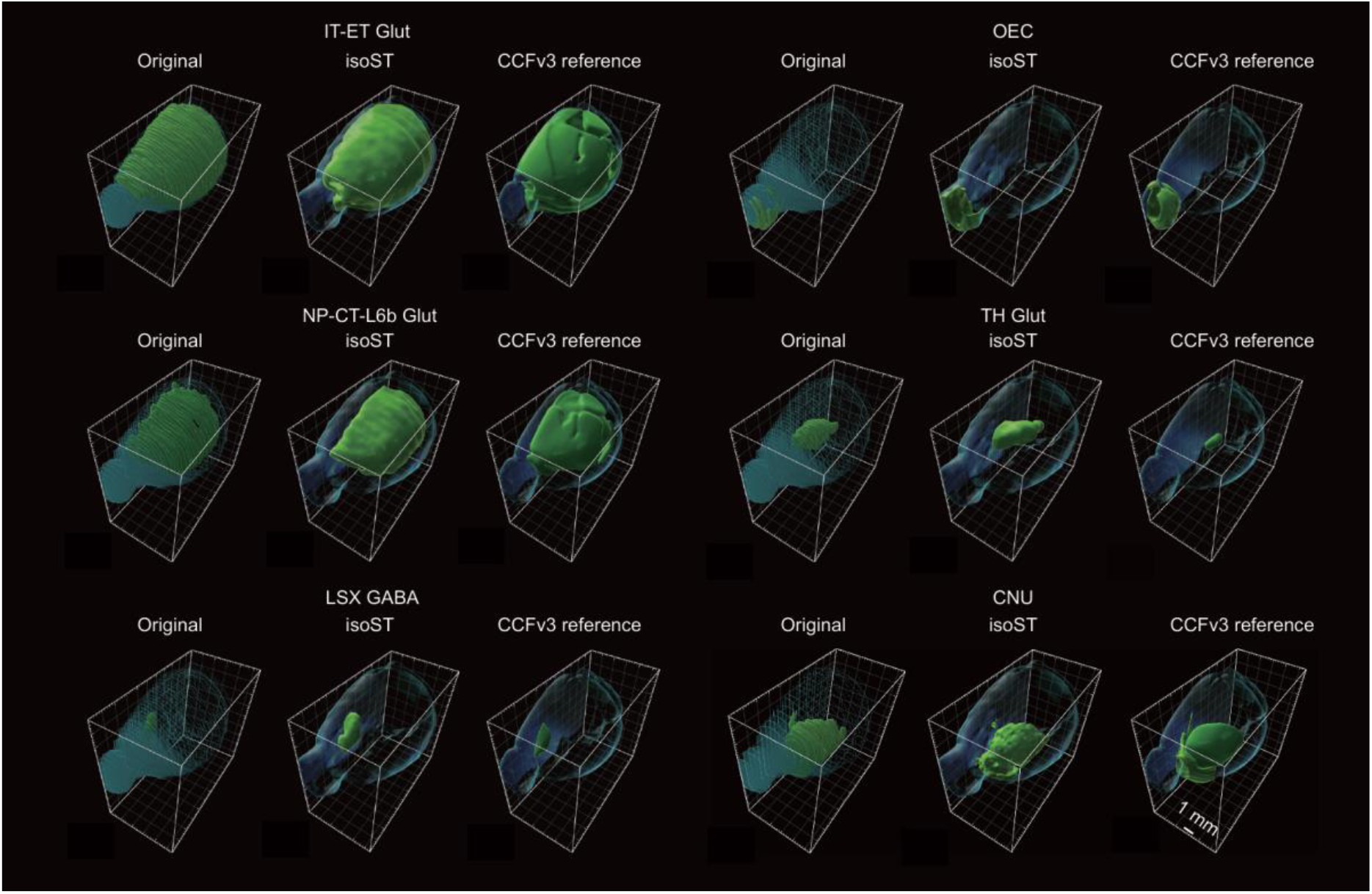
Comparison of original, reconstructed, and CCFv3 reference structures for six representative brain regions. 3D spatial distributions of representative brain regions rendered using IMARIS, shown for original MERFISH slices (left), isoST reconstructions (middle), and CCFv3 annotations (right). The regions include: IT-ET Glut: intratelencephalic-extratelencephalic glutamatergic neurons, OEC: olfactory ensheathing cells, NP-CT-L6b Glut: near-projecting corticothalamic layer 6b glutamatergic neurons, TH Glut: thalamic glutamatergic neurons, LSX GABA: lateral septal complex GABAergic neurons, and CNU: cerebral nucleus neurons.

**Supplementary Figure 5.**
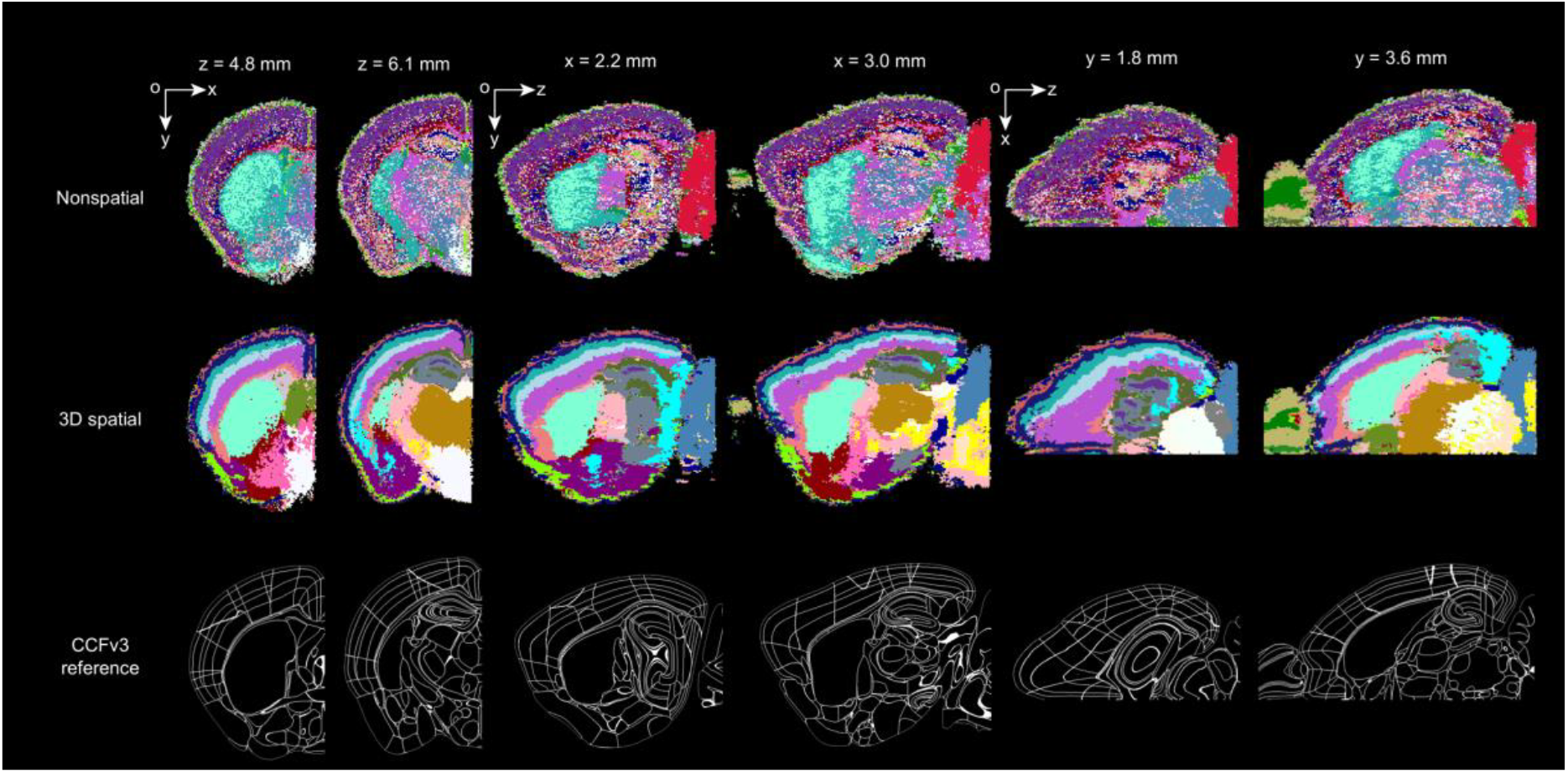
Comparison of Nonspatial and 3D spatial clustering against anatomical references. Representative coronal (z = 4.8 mm, z = 6.1 mm), sagittal (x = 2.2 mm, x = 3.0 mm), and horizontal (y = 1.8 mm, y = 3.6 mm) sections are visualized. Top row: unsupervised clustering results directly obtained from the isoST-reconstructed 3D volume, without additional feature processing. Middle row: clustering results based on 3D spatial features extracted from the same isoST-reconstructed volume. Bottom row: Corresponding anatomical references from the Allen CCFv3 atlas.

**Supplementary Figure 6.**
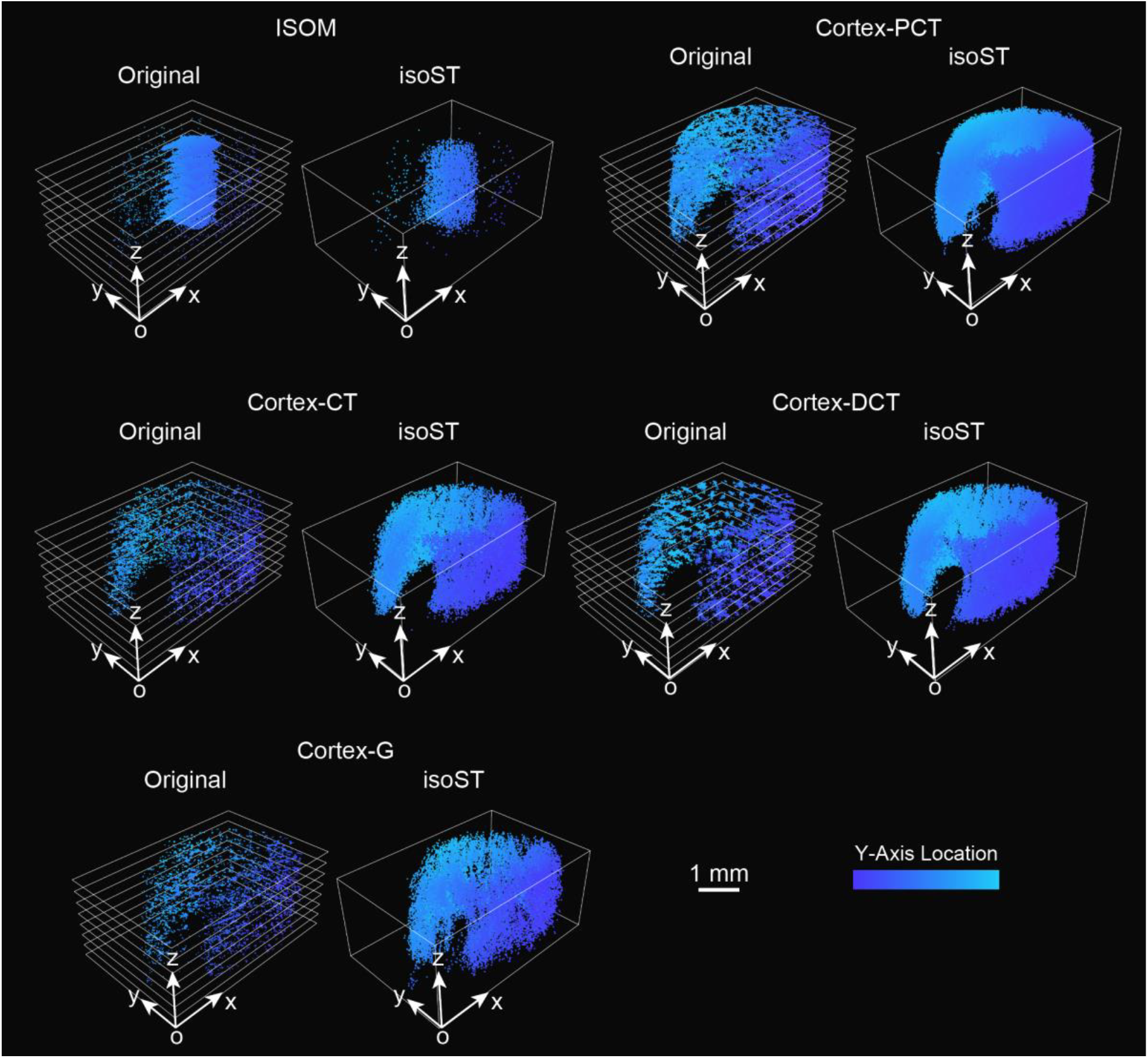
Visualization of kidney tissue structures before and after 3D reconstruction by isoST. 3D spatial distributions of kidney regions. Each structure is visualized in 3D with Y-axis color coding representing vertical anatomical positioning. ISOM: inner stripe of outer medulla, cortex-CT: connecting tubule, cortex-DCT: distal convoluted tubule, cortex-G: glomerulus, and cortex-PCT: proximal convoluted tubule.

**Supplementary Figure 7.**
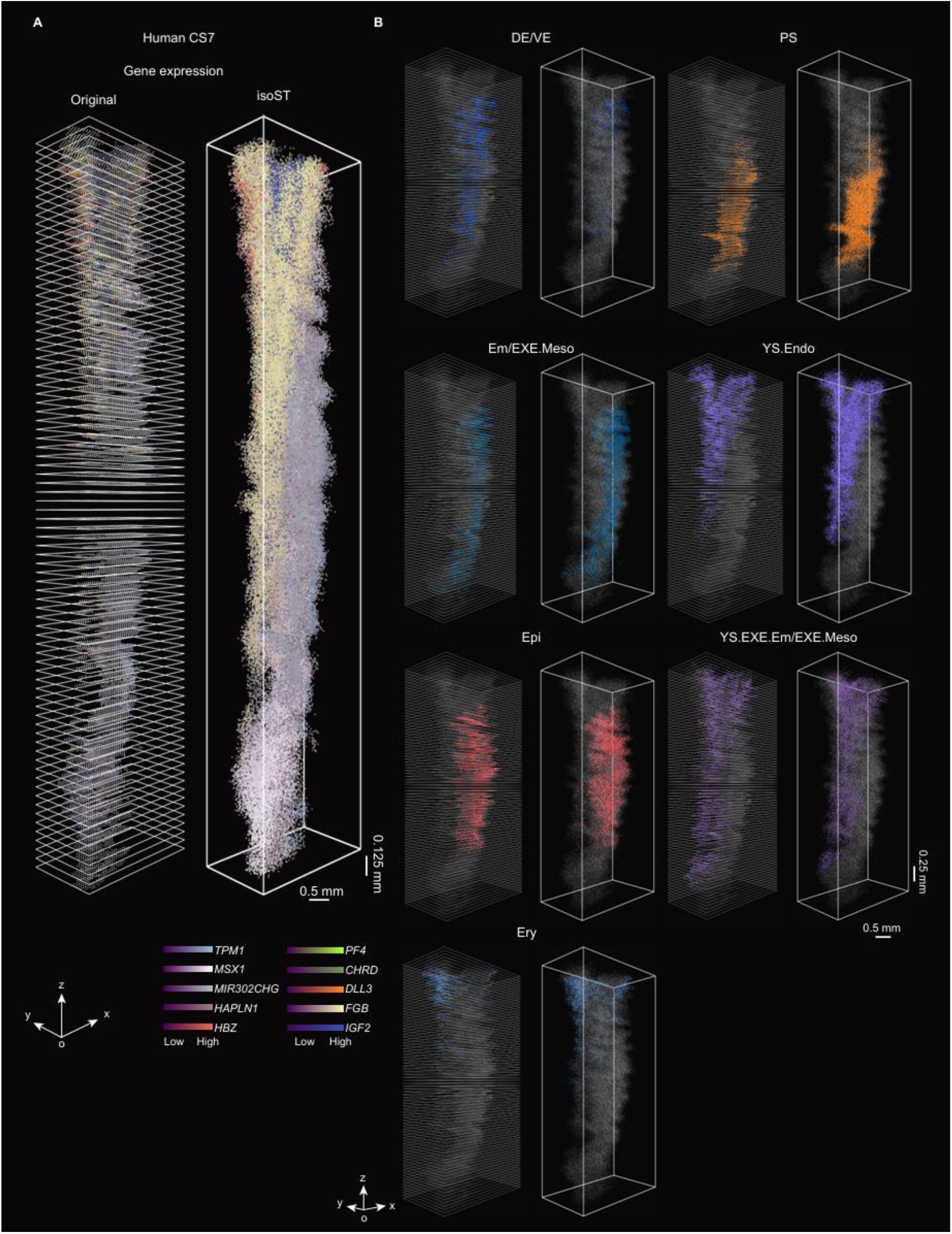
3D reconstruction of the CS7 human embryo using isoST. (**A**) Original 2D spatial transcriptomics sections (*n* = 82) and the corresponding isoST-reconstructed 3D transcriptomic volume for the Carnegie Stage 7 (CS7) human embryo. Expression patterns of the top 10 highly variable genes are shown across the reconstructed volume. (**B**) Spatial distribution of major tissue regions and cell types. DE/VE: definitive/visceral endoderm, PS: primitive streak, Em/EXE.Meso: embryonic/extra-embryonic mesoderm, YS.Endo: yolk sac endoderm, Epi: epiblast, YS.EXE.Meso: yolk sac extra-embryonic mesoderm, and Ery: erythroblast.

**Supplementary Figure 8.**
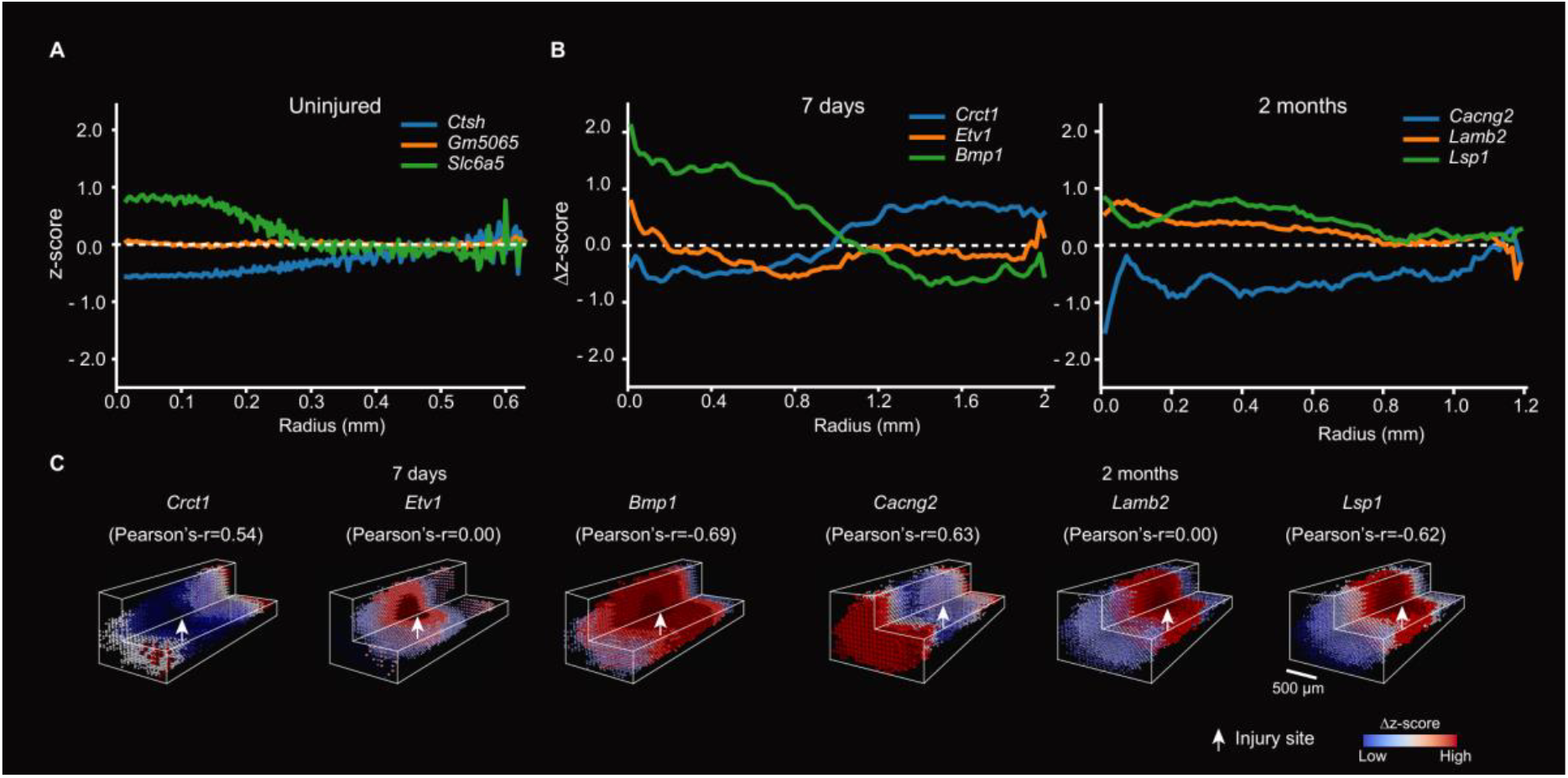
3D gradient analysis of isoST-reconstructed mouse spinal cords. (**A**) Z-scored gene expression profiles plotted against radial distance in uninjured samples, modeled using cylindrical coordinates centered on the spinal canal. (**B**) Δ z-scores of gene expression (post-injury minus uninjured), plotted against spherical distance from the lesion site for samples collected at 7 days and 2 months post-injury.. (**C**) Δz-score patterns of representative genes showing the strongest positive, neutral, and negative spatial correlations, illustrating distinct injury- and recovery-associated trends.

**Supplementary Figure 9.**
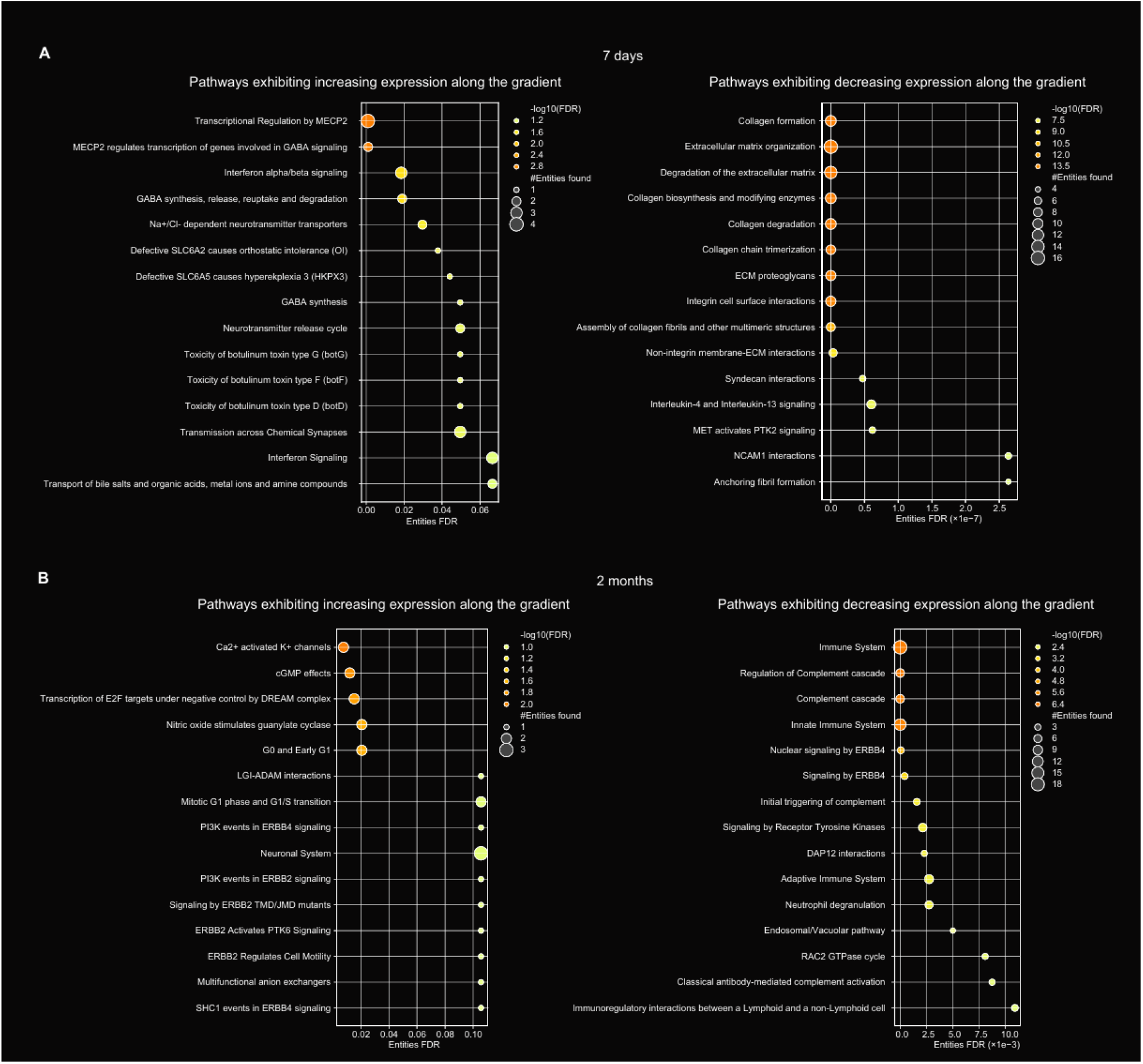
Pathway enrichment analysis along the spherical gradient. Pathways enriched among genes exhibiting significantly increased (left) or decreased (right) expression along the spherical injury gradient at (**A**) 7 days and (**B**) 2 months post-injury. For each time point, the top 15 genes with the most positive and most negative correlations were selected for enrichment analysis. Dot size indicates the number of genes associated with each pathway, and color represents statistical significance (−log_10_(FDR)).

**Supplementary Figure 10.**
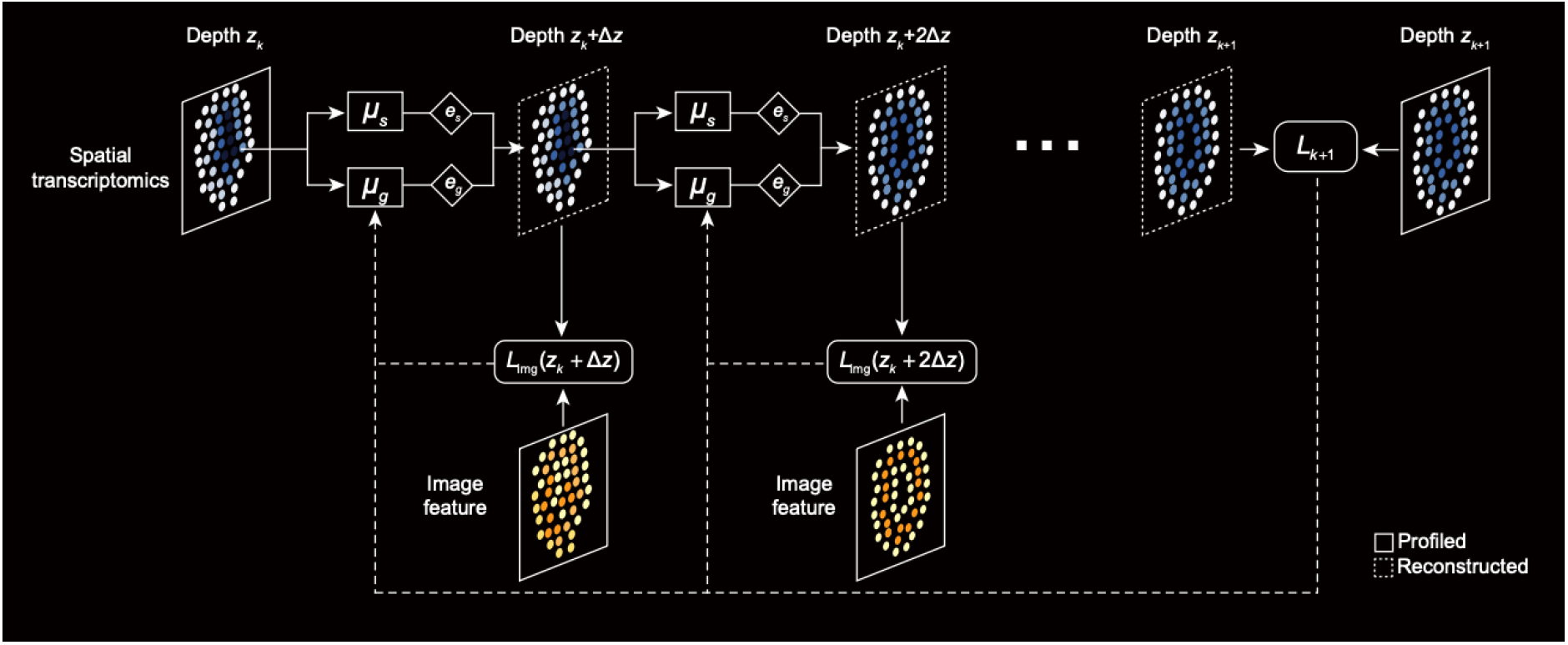
isoST-i framework with image regulations. isoST-i reconstructs intermediate slices between observed depths *z*_*k*_ and *z*_*k*+1_ by iteratively applying learned shape and expression gradient modules ***μ***_s_ and ***μ***_g_. When histological images are available at an imputed depth, an image-based regularization loss *L*_img_ is included to provide additional constraints.

**Supplementary Figure 11.**
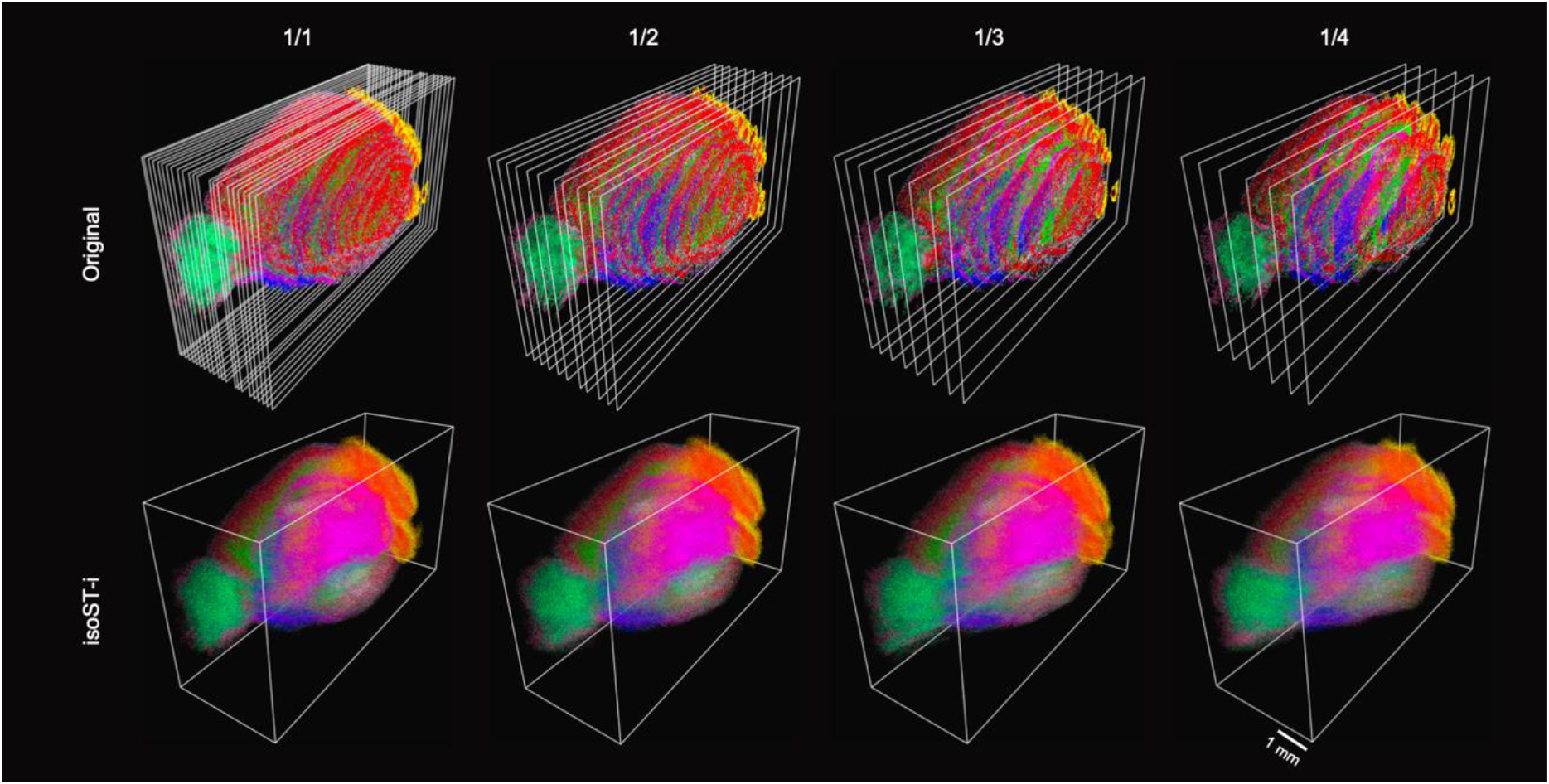
3D reconstruction results using isoST-i under different transcriptomic sampling densities.

**Supplementary Figure 12.**
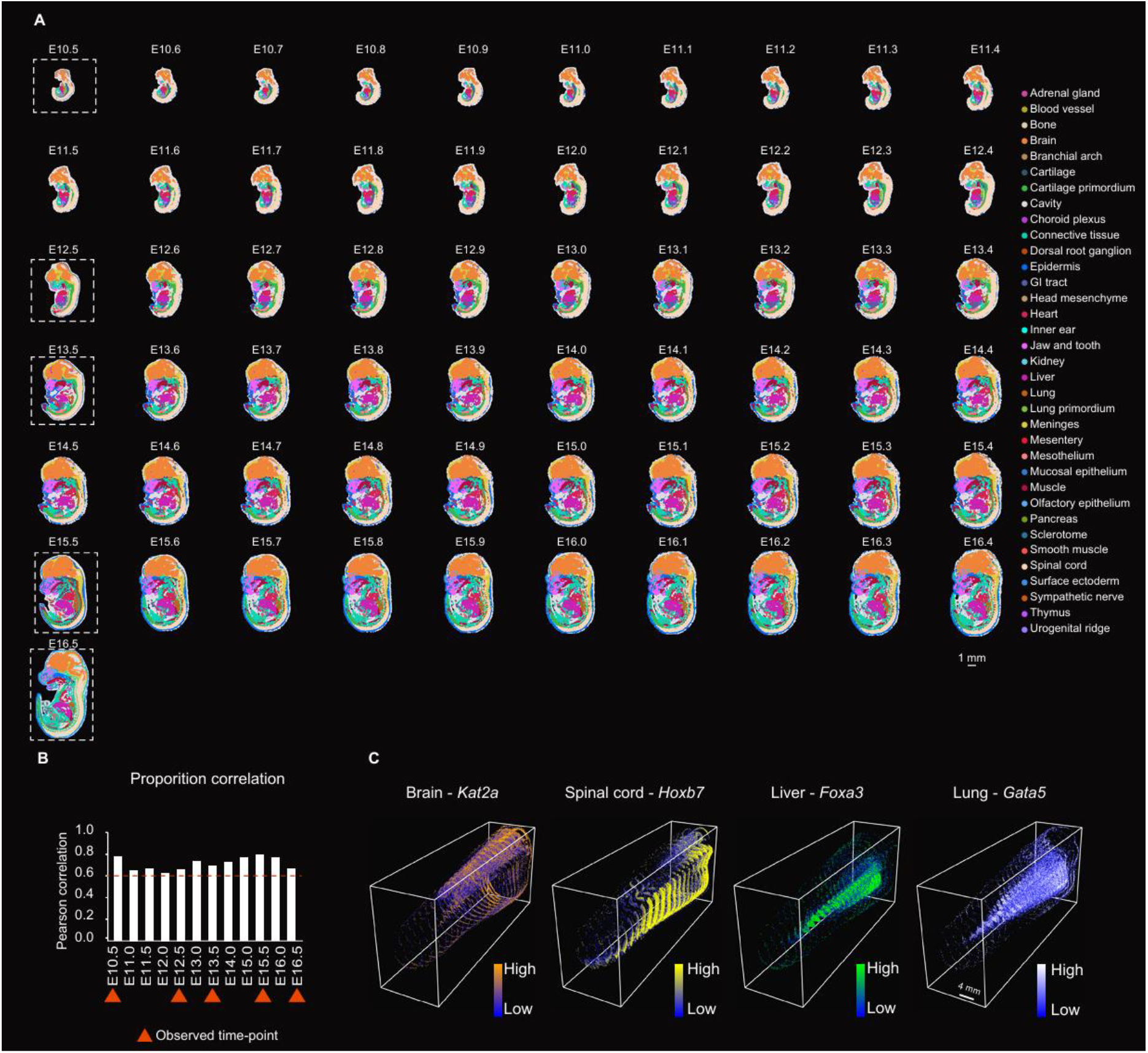
Reconstruction of mouse embryonic development using isoST-t. (**A**) Reconstructed spatial maps of mouse embryos from E10.5 to E16.5 generated by isoST-t based on five observed time points (outlined slices; time points marked by triangles in b). Slices are color-coded by organ annotations. (**B**) Pearson correlation between organ proportions in isoST-t reconstructions and those from a single-cell reference atlas across 12 developmental stages. Orange triangles indicate the five time points used in reconstruction. (**C**) Temporal gene expression dynamics of key developmental regulators across organs: *Kat2a* (brain), *Hoxb7* (spinal cord), *Foxa3* (liver), and *Gata5* (lung). Color scale reflects expression intensity.

## Supplementary Note

**Supplementary Note 1** | **Backward progression of isoST**.

To model the backward progression of isoST, we introduce a set of backward stochastic differential equations (SDEs):

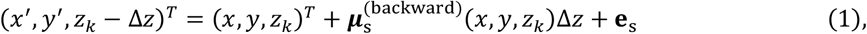

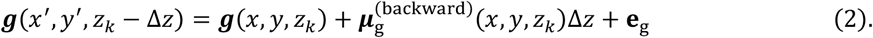

Here, the drift coefficients 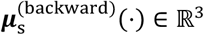 and 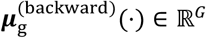 are also learnable functions that model the spatial gradients of tissue morphology and gene expression, respectively, along the reverse direction of the depth axis. Due to the presence of diffusion terms **e**_s_ ∈ ℝ^3^ and **e**_g_ ∈ ℝ^*G*^, these backward drift functions are not simply the negation of their forward counterparts ***μ***_s_ and ***μ***_g_.

To explicitly capture backward dynamics, we parameterize 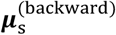 and 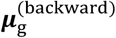 using a separate pair of graph neural networks (GNNs) *h*_*s*_ and *h*_*g*_. The networks *h*_*s*_ and *h*_*g*_ adopt the same architectural designs as their forward counterparts *f*_s_ and *f*_g_, respectively.

## Supplementary Table

**Supplementary Table 1.** isoST hyperparameters across datasets.

| Dataset | $\sigma_{x,y}$ | $\sigma_z$ | $\sigma_g$ | $\Delta z$ | $\alpha$ | Epoch |
| --- | --- | --- | --- | --- | --- | --- |
| Mouse brain | 0.01 | 0.1 | 0.1 | 0.01 | 0.1 | (100, 100, 100) |
| Mouse kidney | 0.01 | 0.14 | 0.1 | 0.1 | 1 | (100, 100, 100) |
| Human embryo | 0.014 | 0.1 | 0.1 | 0.1 | 1 | (100, 100, 100) |
| Mouse spinal cord (uninjured) | 0.01 | 0.1 | 0.1 | 0.01 | 0.1 | (100, 100, 100) |
| Mouse spinal cord (7 days) | 0.014 | 0.14 | 0.1 | 0.02 | 0.02 | (100, 100, 100) |
| Mouse spinal cord (2 months) | 0.01 | 0.1 | 0.1 | 0.01 | 0.1 | (100, 100, 100) |
| Mouse embryo | 0.014 | 0.1 | 0.1 | 0.1 | 0.1 | (2000, 2000, 2000) |

**Supplementary Table 2.** isoST-i hyperparameters under varying input sparsity.

| Reduction rate | $\sigma_{x,y}$ | $\sigma_z$ | $\sigma_g$ | $\Delta z$ | $\alpha$ | $\beta$ | $\gamma$ | Epoch |
| --- | --- | --- | --- | --- | --- | --- | --- | --- |
| 1/1 | 0.01 | 0.1 | 0.1 | 0.01 | 0.1 | 1000 | 0.0001 | (50, 50, 50) |
| 1/2 | 0.01 | 0.07 | 0.1 | 0.01 | 0.1 | 100 | 0.005 | (50, 50, 50) |
| 1/3 | 0.01 | 0.057 | 0.1 | 0.01 | 0.1 | 1000 | 0.0001 | (50, 50, 50) |
| 1/4 | 0.01 | 0.05 | 0.1 | 0.01 | 0.1 | 100 | 0.005 | (50, 50, 50) |

**Supplementary Table 3. Mapping table between cell types and organs** See cell_type_to_organ_mapping.csv

